# Optimizing marker density for maximizing the accuracy of genomic prediction and heritability estimates in three major North American and European spruce species

**DOI:** 10.1101/2025.11.28.691168

**Authors:** André Soro, Patrick Lenz, Jean Beaulieu, Jean-Philippe Laverdière, Simon Nadeau, France Gagnon, Funda Ogut, Harry X. Wu, Martin Perron, Jean Bousquet

## Abstract

Genomic prediction, also called genomic selection (GS), is being increasingly used in tree breeding with aim to accelerate genetic gains by shortening the long breeding cycles. However, high genotyping costs remain a challenge. This study aimed to determine the optimal marker density in genome coverage, to maximize GS accuracy and precision of heritability estimates for growth and wood quality traits. Thousands of SNPs representative of the exome of three major spruce species were used: 18,275 SNPs for black spruce (representing 10,894 distinct gene loci), 11,328 SNPs for white spruce (8647 gene loci), and 116,765 SNPs for Norway spruce (20,695 gene loci). For each species, a similar experimental design was used with related full-sib families replicated on two sites, and GBLUP prediction models were developed. The effect of varying the number of SNPs was examined by re-sampling subsets from 500 to 100,000 SNPs. Results indicated that plateaus in heritability estimates were reached as the marker density increased, stabilizing between 4000 to 8000 SNPs for a spruce genome size of around 2000 centimogans, a trend consistent across all traits and species. Predictive ability and prediction accuracy both increased with the number of SNPs up to a similar level, beyond which further improvements were marginal. Such optimal marker density should be financially attainable for most spruce breeding programs, striking a balance between the need for maximizing accuracy and that for minimizing genotyping costs. These findings should support the further deployment of GS in conifer breeding programs, with high selection precision and by reducing the financial burden of very high-density SNP coverage, even for conifers characterized by large giga-genomes.

## 1 Introduction

Genomic prediction, also known as genomic selection (GS), is a predictive method of selection that relies on dense genome-wide marker coverage to model the full spectrum of quantitative trait loci (QTL) effects across the genome (Meuwissen et al. 2001). But it also depends on the relatedness of individuals making up the training and target populations (Habier et al. 2007; Zapata-Valenzuela et al. 2012; Beaulieu et al. 2014b; Lenz et al. 2017). It has been applied in numerous animal (Meuwissen et al. 2016) and plant breeding programs (Crossa et al. 2017), and is being implemented progressively more into tree breeding programs (Bousquet et al. 2021; Beaulieu et al. 2022, 2024; Grattapaglia et al. 2022).

Conventional breeding methods applied to forest trees, usually characterized by lengthy breeding cycles and large efforts, particularly for conifers (Mullin et al. 2011), have often struggled to keep pace with the urgent need for improved tree varieties that can adapt to rapidly changing market and environmental conditions, especially with the intensification of climate change effects (Depardieu et al. 2020; Benomar et al. 2022; Laverdière et al. 2022). GS has the potential to help boost genetic gain per time unit by reducing the need for lengthy field evaluation of phenotypes as in conventional programs (Grattapaglia and Resende 2011; Beaulieu et al. 2014a,b, 2020; Lenz et al. 2017, 2020a,b; Chen et al. 2018), while increasing selection intensity by screening thousands of candidates and facilitating multi-trait selection addressing multiple breeding objectives (Lenz et al. 2020a; Bousquet et al. 2021). GS offers consequently significant advantages over conventional breeding strategies, which are often time-consuming and costly, particularly for long-lived species such as spruces (Beaulieu et al. 2014a; Park et al. 2016; Lenz et al. 2017, 2020a,b, Lebedev et al. 2020; Bousquet et al. 2021). GS is thus positioned as a crucial tool to help tree breeders meeting the rising demand for high-quality wood fibers for several uses worldwide, but also facing the growing challenges imposed by rapid climate and environmental changes to more rapidly select and produce adapted seedling stocks (Park et al. 2016, Beaulieu et al. 2020, Laverdière et al. 2022).

GS leverages the vast amount of genetic information made available through high-throughput genotyping technologies to make accurate predictions of individuals’ genetic potential for valued quantitative traits (Meuwissen et al. 2001). This is even more true when predictions are made for untested crosses in a tree breeding context as genomic markers can better trace non-additive effects such as dominance (Nadeau et al. 2023). Growth traits which are widely used in tree breeding strategies, are generally under lower additive genetic control and often carry important dominance components whereas wood quality traits are mostly under moderate to strong additive genetic control (Lenz et al. 2020). Thus, GS has the potential to be efficient to select for traits of various genetic architecture (Grattapaglia and Resende 2011) and to accelerate the identification of superior individuals for reforestation based on their growth, wood quality, and resistance to biotic or abiotic stressors (Beaulieu et al. 2020; Lenz et al. 2020a; Laverdière et al. 2022). Most economically important traits are of quantitative nature and controlled by many genes having each a small effect on the phenotype (White et al. 2007). Therefore, for GS to be successful, it is essential that an adequate number of markers is used to achieve a sufficient genome coverage and that at least one of these markers is in more or less complete linkage disequilibrium (LD) with each of the QTLs that affect the quantitative trait of interest (Meuwissen et al. 2001; Hayes and Goddard 2010).

Early GS applications in forest trees were bolstered by advancements in genotyping technologies, statistics and information technology (IT). High-throughput genotyping technologies were indeed made available early to forest geneticists (e.g. Pavy et al. 2008, 2013). Various statistical methods such as ridge regression, Best Linear Unbiased Prediction (BLUP), especially applied to genomic data (GBLUP) as well as Bayesian methods, allowed for more accurate genetic predictions by effectively incorporating dense marker information and accounting for the complex genetic architecture of quantitative traits (De Los Campos et al. 2013). Algorithms and softwares were rapidly developed by statisticians under the R Project (R Core Team 2023) and made available to the genetics community. Grattapaglia and Resende (2011) demonstrated the potential of GS to improve traits with moderate to high heritability in *Eucalyptus* and through simulations, setting the baseline for similar approaches in other broadleaves and conifer species.

In recent years, significant progress has been made in the publication of proof of concept studies of GS on conifers, especially for species with advanced genetic improvement program underway (Mullin et al. 2011), such as black spruce (*Picea mariana* [Mill.] B.S.P.) (Lenz et al. 2017), white spruce (*Picea glauca* [Moench] Voss) (Beaulieu et al. 2014a,b; Ratcliffe et al. 2017; Beaulieu et al. 2020; Laverdière et al. 2022; Nadeau et al. 2023), Norway spruce (*Picea abies* [L.] Karst.) (Chen et al. 2018, 2019; Lenz et al. 2020a), loblolly pine (*Pinus taeda* L.) (Resende et al. 2012; Zapata-Valenzuela et al. 2013), maritime pine (*Pinus pinaster* Aiton) (Bartholomé et al. 2016, Monterey pine (*Pinus radiata* D. Don) (Li and Dungey 2018; Freeman et al. 2022), lodgepole pine (*Pinus contorta* Douglas) (Ukrainetz NK, Mansfield SD 2020), Douglas-Fir (*Pseudotsuga menziesii* Mirb. [Franco]) (Thistlethwaite et al. 2017), japanese cedar (*Cryptomeria japonica* D. Don) (Hiraoka et al. 2018), western red cedar (*Thuja plicata* Donn ex Don) (Gamal El-Dien et al. 2022, 2024). These studies relied on SNP markers to capture the genetic variation within species, facilitating the accurate prediction of complex traits such as growth rate, wood quality, and pest resistance or tolerance.

Despite the technical progress, several challenges remain in the implementation of GS in forest tree breeding. One of the primary issues is to increase marker density to improve capture of short-range LD, whilst determining the optimal number of markers required for accurate genomic predictions. For across-generation GS accuracy in advanced-breeding programs, increasing marker density might even be more important to capture LD. While high-density SNP arrays might provide more precise genetic information, they are often cost-prohibitive for large-scale tree breeding programs. However, research by Habier et al. (2007) highlighted the trade-offs between marker density, cost, and prediction accuracy, emphasizing the need for cost-effective strategies that do not compromise the reliability of genetic predictions.

Simulations demonstrated early on that both marker density and breeding population status number (or effective population size) have important impacts on the accuracy of GS (Grattapaglia and Resende 2011). Using a meta-analysis of GS studies in tree species, Beaulieu et al. (2024) confirmed that the higher the breeding population status number, the lower the GS accuracy for a marker density of similar magnitude. They also showed that in the range of numbers of markers used in published studies of GS in forest tree species, there was no significant variation in GS model accuracy. In contrast to tree breeding, genomic selection (GS) is more advanced in plant and animal breeding (Schaeffer 2006; Hayes et al. 2009). In these fields, several studies investigated the effect of marker density on prediction accuracy, including in wheat (Norman et al. 2018), maize (Guo et al. 2020), alfalfa (Wang et al. 2024), cattle and pig (Su et al. 2012; Zhang et al. 2019; Reverter et al. 2020). For instance, Wang et al. (2024) showed in alfafa that GS prediction accuracy reached a plateau with 3000 SNPs, and with around 5000 SNPs for wheat and maize (Norman et al. 2018; Guo et al. 2020), despite their large differences in physical genome size.

Several authors also reported for tree species that both heritability and prediction accuracy estimated by GS appear to level out at quite low marker density (Lenz et al. 2017; Tan et al. 2017; Chen et al. 2018; Calleja-Rodriguez et al. 2020; Estopa et al. 2023). Hence, it has been hypothesized that only a relatively modest number of markers is needed to capture genetic relatedness in structured breeding populations because prediction accuracy does not appear to be largely controlled by marker-QTL LD but mostly by relatedness in such populations (Habier et al. 2007; Zapata-Valenzuela et al. 2012; Beaulieu et al. 2014b; Lenz et al. 2017; Grattapaglia et al. 2022). However, previous studies have not consistently established the optimal marker density required to reach most accurate GS predictions and precise estimation of genetic parameters, though the level of marker density necessary for this appears to be quite coherent across published studies on tree species (Beaulieu et al. 2024), and in agreement with early simulation results (Grattapaglia and Resende 2011) indicating that relatively low marker densities should be sufficient to attain high GS accuracy.

The various proof of concept studies on the application of GS in spruce species, during the last decade, have relied on a quite variable number of SNPs ranging from about 5000 to 100,000, with most of them being at the lower end of this range (see Beaulieu et al. [2024] for an exhaustive list). For example, Nadeau et al. (2023) used 4092 SNPs from distinct gene loci in their GS study on white spruce whereas Chen et al. (2018) used 116,765 SNPs with quite high redundancy of SNPs per gene locus for their application of GS in Norway spruce. Further tests on the effect of the number of markers used in GS on the heritability estimates obtained and on the prediction accuracy would allow deepening knowledge and contribute to the application of GS to tree breeding programs.

In the current study, data from three major spruce species in the northern hemisphere were assembled. Black spruce is one of the most widely distributed and resilient conifers in North America’s boreal forests. It is a species of great importance in Canada and the northern United States, particularly for the production of high-quality pulp and solid wood products, especially framing materials (Zhang and Koubaa 2008). As for many other commercially valuable species, it is a key focus for breeding programs aimed at improving various growth and wood quality traits (Mullin et al. 2011), and is widely planted in Ontario and Quebec (Natural Resources Canada 2024).

Of a similar geographic range, white spruce is another commercially significant species in the boreal forests of North America. It is reproductively isolated and quite phylogenetically remote from black spruce (Bouillé et al. 2011) and is also widely utilised in the production of pulpwood and solid wood products (Zhang and Koubaa 2008), and the subject of many breeding programs (Mullin et al. 2011). Owing to its high survival rate, adaptability to diverse ecological conditions, and rapid growth, it is one of the most productive conifers in boreal and temperate forests in Canada and is commonly planted, particularly in Alberta, Quebec, and the Maritime provinces.

Norway spruce is native to Scandinavia, Eastern Europe and the mountainous regions of Western and Central Europe, and it represents the most important conifer for lumber and pulp and paper production in Europe (Hannrup et al. 2004; Mullin et al. 2011). Indeed, it contributes to 40% of the total growth in Europe Nordic forests, establishing it as a highly important commercial tree species and the subject of many tree breeding programs (Mullin et al. 2011). It has also been introduced in eastern Canada, where it is one of the most productive conifers in plantations (Lenz et al. 2020a). Breeding programs have hence been developed for this species to enhance various traits, including climate change adaptation, wood quality and eastern white pine weevil resistance. Altogether, the economic and ecological importance of these three species justify the quest for more efficient breeding methods and make them good candidates for the operationalization of GS in tree breeding efforts.

Based on extensive datasets drawn from operational breeding trials for these three widely reforested spruce species of the northern hemisphere, the present study aims to determine the optimal marker density that maximizes GS prediction accuracy and the precision of genetic parameter estimates for traits with contrasted heritability in spruce breeding populations. Using the same GS modeling approach for the three species, our objective was to determine at which level of marker density, estimates of heritability and GS prediction accuracy for both growth and wood quality traits reached plateaus of values. In short, these plateaus were attained with less than 10,000 markers, beyond which further improvements were marginal and not significant.

## 2 Materials and methods

### 2.1 Sampling of plant material

For this study, three major datasets respectively for black, white and Norway spruces were obtained. The black spruce dataset was collected in a 33-year-old progeny trial previously used in a previous GS study (Lenz et al. 2017) and replicated on two sites, Matapedia (48° 32’N, 67° 25’W, altitude: 216 m) and Robidoux (latitude: 48° 18’N, longitude: 65° 31’W, altitude: 275 m). The 918 sampled individuals belonged to 34 controlled-pollinated families derived from a partial diallel mating design. The 46 parents originated from 9 provenances from the Canadian provinces of Ontario, New Brunswick and Manitoba, and one from Maine in the northeastern United States. Progeny were raised as rooted cuttings in a nursery for three years and then established in 1991 on two forest sites using a randomized complete block design, with 4-tree row plots and a 2 m by 2 m spacing.

For the white spruce dataset, 988 18-year-old trees were sampled from 56 full-sib families originating for 39 parents arranged in a partial diallel, corresponding to a large subset of the trees analysed by Nadeau et al. (2023). In this study, we used material from two of the six breeding groups, namely the breeding groups 3 and 4 as labelled in Nadeau et al. (2023). The progeny test was established on two contrasting sites, Asselin (47° 55’ N, 68° 26’ W, altitude: 278 m) and St. Casimir (46° 42’ N,72° 06’ W, altitude: 52 m). The experimental layout followed a randomized complete block design with 10 replications. Trees were planted in row-plots, with each plot consisting of five trees spaced 2 m by 2 m apart.

For Norway spruce, a dataset previously published by (Chen et al. (2018) was used. Briefly, 1370 individuals were selected from two 28-year-old control-pollinated progeny trials replicating the same 128 full-sib families from a partial diallel mating design that consisted of 55 parents originating from Northern Sweden. Five to 20 trees per family per site were selected from the Vindeln (64.30°N, 19.67°E, altitude: 325 m) and Hädanberg (63.58°N, 18.19°E, altitude, 240 m) testing sites of same age. A completely randomized design without designed pre-block was used in the Vindeln trial (site 1), which was divided into 44 post-blocks. The same design was also used in the Hädanberg trial (site 2) with 44 post-blocks, but also an extra design with 47 extra plots. After a spatial analysis, the 47 plots were combined into two large post-blocks for final analyses (Chen et al. 2018).

### 2.2 Trait phenotyping

A similar set of phenotypic traits was considered for all three species. For black spruce, three phenotypic traits were assessed at 30 years after plantation: tree height, diameter at breast height (DBH, measured at 1.3 meters above ground) and wood density. In 2021, a wood increment core was extracted from the south-facing side of each tree. The cores were stored in a freezer, conditioned to 7% moisture content, and cut to a thickness of 1.68 mm prior to X-ray densitometry analysis using Quintek Measurement Systems (Knoxville, TN). Wood density was calculated as a ring area-weighted mean from the recorded pith-to-bark wood density profiles.

For white spruce, four phenotypic traits were considered: tree height, DBH, wood density, and acoustic velocity. They were assessed at 13 years after plantation. Wood density was determined with X-ray densitometry as described above (Beaulieu et al. 2014b). Acoustic velocity, which is a proxy for wood stiffness (Lenz et al. 2013), was measured on standing trees with the Hitman ST300 tool (Fibre-gen, Christchurch, New Zealand).

For Norway spruce, three phenotypic traits were considered: tree height at the age of 17 years, Pilodyn pin penetration as a surrogate for wood density, and acoustic velocity using the Hitman ST300 tool.

### 2.3 SNP genotyping

For black spruce, 918 trees were genotyped with the new Infinium iSelect SNP array Pmr25k (Illumina, San Diego, CA) (see Gérardi et al. 2025 for details), from which 18,275 SNPs segregated in black spruce, representing 12,438 distinct gene loci. For white spruce, two genotyping datasets were combined for same trees: earlier published genotyping data from the Infinium SNP array PgLM3+ (Pavy et al. 2013, Beaulieu et al. 2014a) with 6932 SNPs representing 6918 distinct gene loci, and the other one from using the Infinium SNP array Pmr25k with 4747 SNPs segregating for white spruce and representing 3991 gene loci, for a total of 11,328 non-redundant SNPs representing a total 8647 distinct gene loci. To retain only high-quality non-singleton SNPs, the following SNP quality criteria were applied: a genotyping reproducibility rate ≥ 95% as calculated from replicated controls, a call rate ≥ 90%, a fixation index |*F*_e_| ≤ 0.50, and a minor allele frequency (MAF) ≥ 0.01. Mendelian segregation of each SNP was also verified. For black spruce, SNPs with ≥ 5% of genotyping errors, that is incompatible genotypes according to parent genotypes, or for which genotype frequencies strongly departed from Mendelian expectations in more than 3 families (Fisher exact test, *P* < 0.001 using families with ≥ 10 individuals) were also discarded. For white spruce, see Nadeau et al. (2023) for details of the filtering criteria including Mendelian segregation, similar to those used for black spruce above. A total of two black spruce trees and one white spruce trees were discarded because their call rates were <90% (average call rate was 99%). After applying these filters for retaining only non-singleton high-quality SNPs, a total of 16,207 SNPs representative 10,894 distinct gene loci were retained for quantitative genomic analyses for black spruce, and a total of 10,289 SNPs representative 8,647 distinct gene loci for quantitative genomic analyses for white spruce. For black spruce and white spruce, the retained SNPs had an average call rate per tree of 99.0%, an average genotyping reproducibility rate of 99.99% as assessed by two replicated control genotypes on each genotyping plate, and an overall average minor allele frequency MAF = 0.19 and an average fixation index *F_e_* = 0.04.

For Norway spruce, the 1375 trees were genotyped for 116,765 SNPs representing 20,695 distinct gene clusters, using an exome capture and Illumina HiSeq 2500 platform (Chen et al. 2018). After applying several filters for retaining high-quality SNPs with a minor allele frequency MAF ≥ 0.01 with call rate ≥ 90% (116,219 SNPs were retained for analyses, as in Chen et al. (2018).

Overall, no imputation of missing data was necessary given the high quality of the genotyping data sets obtained for each of the three species, with individual trees having call rate in excess of 99%, on average.

### 2.4 Quantitative genetic analyses

#### 2.4.1 Relationship matrices

To estimate genetic parameters for broad- and narrow-sense heritability, individual-tree linear mixed models using pedigree-based or genomic-based relationship matrices for ABLUP-AD and GBLUP-AD methods, respectively were run. The pedigree-based additive relationship matrix (***A***) and its inverse were computed for each species separately using the “Amatrix” function of the R package AGHmatrix (Amadeu et al. 2016) and the “G.inverse” function of the R package ASRgenomics (Gezan et al. 2022), respectively. The realized additive genomic relationship matrix (***G***_***a***_) was obtained using the “G.matrix” function of the R package ASRgenomics, using the formula described by VanRaden (2008). To facilitate the inversion of ***G***_***a***_, a blending of both matrices was done prior to the inversion of ***G***_***a***_***blended***_such as:

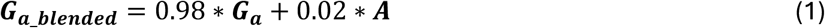

The blending was also tested with proportions of 0.05 and 0.01 for ***A***, but did not change the results of genetic parameters (not shown). The inverse of the matrix ***G***_***a***_***blended***_ was obtained using the “solve” function from base R. Dominance relationship matrices were also calculated. The pedigree-based dominance relationship matrix (***D***) and its inverse were calculated using the “Amatrix” function of the R package AGHmatrix and the “makeD” function of the R package nadiv (Wolak 2012), respectively. The marker-based dominance relationship matrix (***G***_***d***_) was computed using the “G.matrix” function of the R package ASRgenomics following the classical parametrization as in Vitezica et al. (2013), and then blended with the matrix ***D*** as in (1) to invert it with the “G.inverse” function.

#### 2.4.2 Statistical models

Every statistical model fitted in this study was performed under ASReml-R version 4 in the R 4.3.2 environment (R Core Team 2023). We built models for several subsets of SNPs, varying in intervals of 1000 from 500 SNPs up to the entire set of available SNPs for each species. For each SNP subset, we first created five replicates by randomly drawing SNPs from the complete set of SNPs. The variation between replicates was estimated by standard deviations. Given that the black spruce test contained some clonal replicates, only one ramet per clone was randomly chosen for each of the five replicates per level of SNP subsampling. To ensure that five replicates were sufficient per subset of SNPs tested in each species to obtain consistent standard deviation among spruce species, additional resampling of up to 20 replicates was conducted for one growth trait (tree height) and one wood quality trait (wood density) in white spruce and Norway spruce, the species in this study with the lowest and highest numbers of available SNPs, respectively (see supplementary material, Fig. S4 and S5). Cross-validations (see below) were conducted for the various subsets of randomly drawn SNPs, i.e., also varying in 1000 SNP intervals from 500 to the full set of SNPs available for each of the replicate and for each of the three species analysed.

Models included additive (A) and dominance (D) effects. In addition to marker-based models, pedigree-based models were also fitted for comparison (GBLUP-AD and ABLUP-AD, respectively).

The statistical model used for black and white spruce is defined below, was such as:

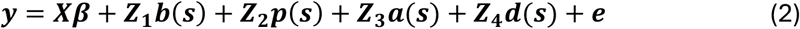

where **d** the vector of measured trait values (phenotypes); ***β*** is the vector of the fixed site effect, including overall mean and the site effect; ***b***(***s***) is the random block within site effect, described as heterogeneous variances between sites using a block diagonal variance-covariance structure, that is 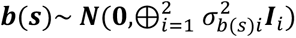; ***p***(***s***) is the random plot effect nested within site, and described as heterogeneous variances between sites using a block diagonal variance-covariance structure, that is 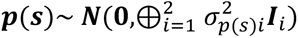; ***a***(***s***) is the random additive effect nested within site described as ***a***(***s***)∼ ***N***(**O**, ***V***_*a*_ ⊗ ***G***_*a*_); ***d***(***s***) is the random dominance effect nested within site described as ***d***(***s***)∼ ***N***(**O**, ***V***_*a*_ ⊗ ***G***_*a*_); and ***e*** is the residual error term described as heterogeneous variances between sites, that is 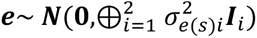. The symbols ⊕ and ⊗ refer to the direct sum and Kronecker product of matrices, respectively.

The statistical model used for Norway spruce is defined below, such as:

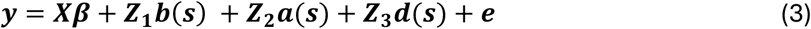

where the terms were previously defined in Eq. (2).

***X*** and ***Z***_***x***_ are incidence matrices of their corresponding effects. The description of the effects fitted in each model is included in Table 1.

**Table 1:**
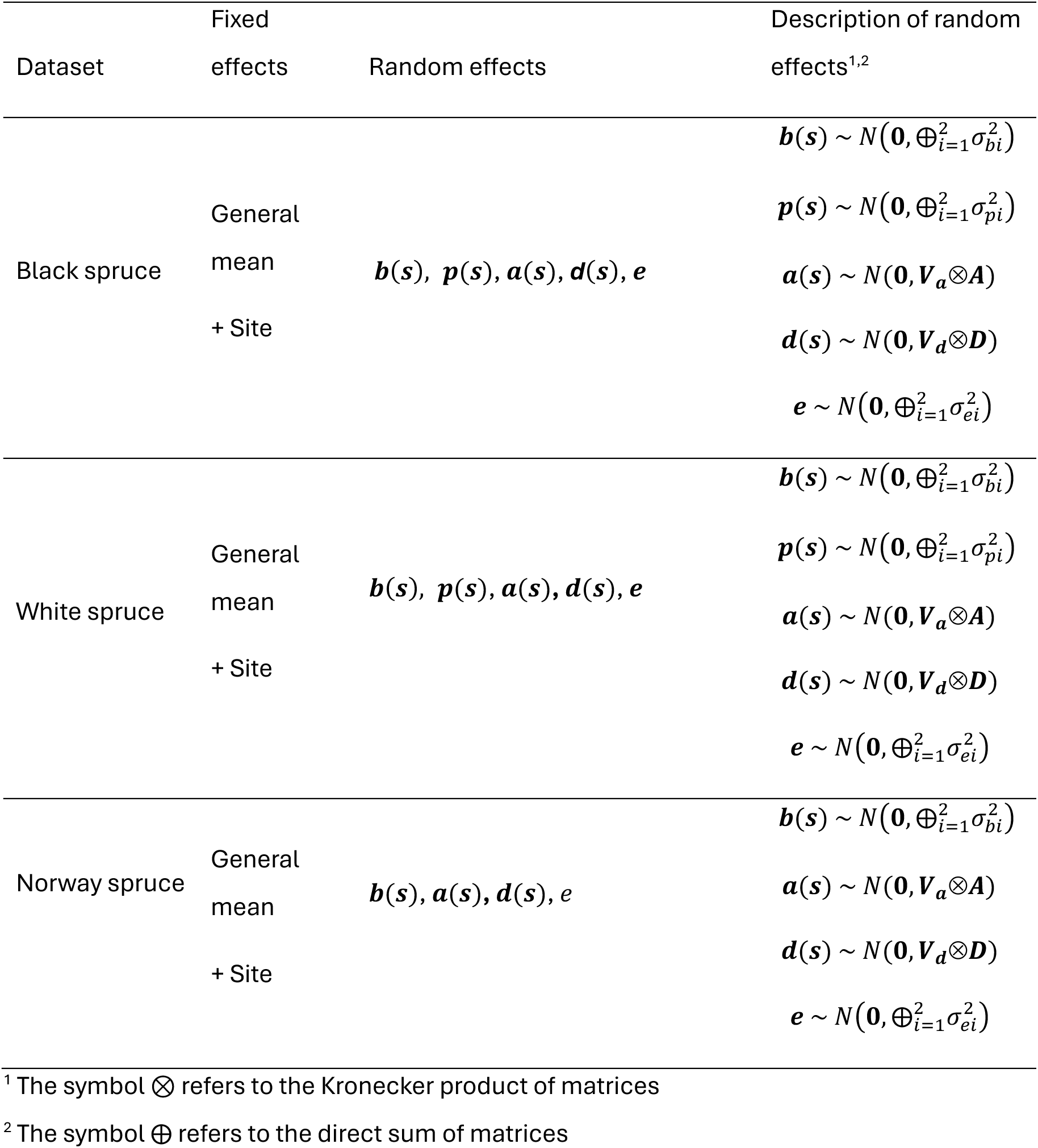
Description of the effects included for each model tested for each dataset.

***A*** and ***D*** matrices for the ABLUP-AD methods were replaced by ***G***_***a***_ and ***G***_***d***_ matrices for the GBLUP-AD method, respectively. The matrices ***V***_***a***_ and ***V***_***d***_ are 2 x 2 variance matrices defined by the correlation of the corresponding genetic effects between sites (*r*_*a*_, *r*_*d*_) and unique variances for each site. They are defined below, such as:

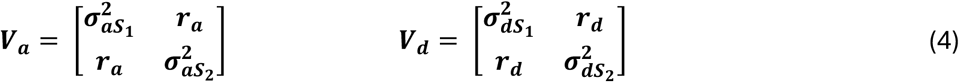

where *σ*^2^_*aS*1_ and *σ*^2^_*aS*2_ are the additive variances of sites 1 and 2, respectively. *σ*^2^_*dS*1_ and *σ*^2^_*dS*2_ are the dominance variances of sites 1 and 2, respectively.

#### 2.4.3 Calculation of broad- and narrow-sense heritability estimates and their components

The across-site additive and dominance genetic variances were recovered as 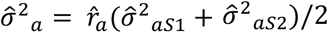 and 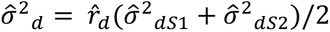, where the subscripts S1 and S2 refer to sites 1 and 2, respectively. The total phenotypic variation 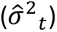 was estimated as the sum of across-site plot, additive, dominance, and residual variances, with 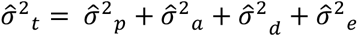. The formulas used for estimations of individual narrow-sense heritability 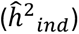, dominance ratio 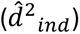, and broad-sense heritability 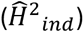 are presented in Eq (5, 6 and 7) below, such as:

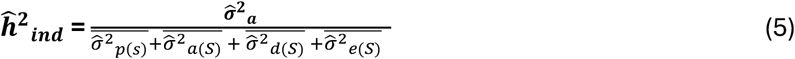

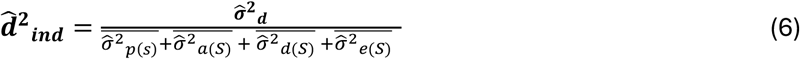

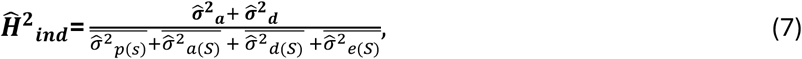

where 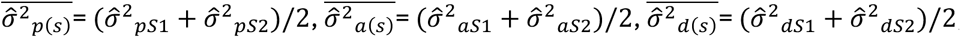, and 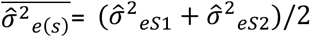.

#### 2.4.4 Cross-validation of genomic selection models

For each dataset analysed, genotypes were randomly split into 10 folds, making sure that each fold contained around 10 percent of the genotypes from each family (within-family folding, Lenz et al., 2020a,b). For the white and Norway spruce datasets, genotypes were always associated with distinct trees. For the black spruce dataset, some genotypes were clonally replicated as indicated above. In this case, one ramet per clone was randomly chosen. For each round of cross-validation, 9 folds were used to train the model used to predict the breeding or genetic values of the remaining fold. Each fold was used once as model validation. From the cross-validation models, the estimated breeding values (EBVs) were recovered from the best linear unbiased predictions (BLUPs) for each genotype associated with the additive relationship matrix. The estimated total genetic values (EGVs) were calculated as the summation, for each genotype, of the additive (EBV) and dominance (EDV) BLUPs. EBVs and EDVs were recovered at the site on which the genotype was in the field. The breeding and genetic values were recovered at the site level. For each repetition, predictive ability (*PA*) and prediction accuracy (*PACC*) were recovered by using the predicted values of the genotypes from all the folds and calculating the correlations within each site separately (i.e. using the breeding/genetic value from the site where the genotyped tree was measured). Then, the *PA* and *PACC* were averaged over sites and repetitions.

The predictive ability was calculated as the Pearson correlation coefficient between the predicted breeding values (*PA*_*BV*_; for the models with additive or additive and dominance effects) or predicted genetic values (*PA*_*GV*_; for the models with additive and dominance effects only) of the validation trees and the observed phenotypes.

The prediction accuracy of breeding values (*PACC*_*BV*_) was estimated as the correlation between estimated breeding values and true breeding values. The prediction accuracy of genetic values (*PACC*_*GV*_) was estimated as the correlation between estimated genetic values and true genetic values. The true narrow-sense and broad-sense heritability estimates were recovered from the full models and calculated for the respective site in the *PA* and *PACC* calculations.

## 3 Results

With the aim to compare trends in heritability estimates and GS model accuracy across all species and trait combinations, we retained very similar models for analyses and comparisons. Initial results had revealed significant variation in site-specific estimates, highlighting the necessity of fitting heterogeneous variances across environments on the different design and genetic effects (Table 2 and S1 to S3). For white and Norway spruces, wood quality traits, namely acoustic velocity and wood density, were moderately to highly heritable at the narrow-sense and broad-sense levels, and this either for the pedigree-based ABLUP-AD or the marker-based GBLUP-AD method; whereas for growth traits, namely tree height and DBH, they were under lower to moderate genetic control (Table 2 and S1 to S3). However, the trend was slightly different in black spruce where all three wood and growth traits were under moderate to high genetic control. Overall, generally higher broad-sense and narrow-sense heritability estimates were observed for the black spruce dataset compared with the Norway spruce and white spruce datasets.

**Table 2:** Across-site genetic parameters estimated using the ABLUP-AD or the GBLUP-AD model including all SNPs available for each species. DBH is tree diameter at breast height.

|  | Parameter | Tree height | DBH | Acoustic velocity | Wood density |
| --- | --- | --- | --- | --- | --- |
| ABLUP-AD |  |  |  |  |  |
| Black | $\hat{h}^2_{ind}$ | 0.56 (0.07) | 0.41 (0.10) | - | 0.32 (0.10) |
| spruce | $\hat{d}^2_{ind}$ | 0.00 (0.00) | 0.11 (0.36) | - | 0.11 (0.11) |
| | $\hat{H}^2_{ind}$ | 0.56 (0.07) | 0.52 (0.37) | - | 0.43 (0.10) |
| White | $\hat{h}^2_{ind}$ | 0.14 (0.10) | 0.05 (0.09) | 0.40 (0.10) | 0.34 (0.09) |
| spruce | $\hat{d}^2_{ind}$ | 0.41 (0.20) | 0.32 (0.16) | 0.01 (0.06) | 0.03 (0.07) |
| | $\hat{H}^2_{ind}$ | 0.55 (0.17) | 0.37 (0.13) | 0.42 (0.11) | 0.38 (0.10) |
| Norway | $\hat{h}^2_{ind}$ | 0.07 (0.04) | - | 0.38 (0.09) | 0.36 (0.08) |
| spruce | $\hat{d}^2_{ind}$ | -0.04 (0.01) | - | 0.11 (0.07) | -0.05 (0.08) |
| | $\hat{H}^2_{ind}$ | 0.03 (0.01) | - | 0.50 (0.10) | 0.31 (0.10) |
| GBLUP-AD |  |  |  |  |  |
| Black | $\hat{h}^2_{ind}$ | 0.46 (0.08) | 0.32 (0.07) | - | 0.31 (0.08) |
| spruce | $\hat{d}^2_{ind}$ | 0.00 (0.00) | 0.12 (0.10) | - | 0.11 (0.10) |
| | $\hat{H}^2_{ind}$ | 0.46 (0.12) | 0.43 (0.09) | - | 0.42 (0.09) |
| White | $\hat{h}^2_{ind}$ | 0.08 (0.06) | -0.01 (0.06) | 0.45 (0.05) | 0.31 (0.06) |
| spruce | $\hat{d}^2_{ind}$ | 0.46 (0.16) | 0.42 (0.17) | 0.00 (0.00) | 0.04 (0.07) |
| | $\hat{H}^2_{ind}$ | 0.54 (0.15) | 0.41 (0.15) | 0.45 (0.05) | 0.35 (0.08) |
| Norway | $\hat{h}^2_{ind}$ | 0.05 (0.04) | - | 0.26 (0.07) | 0.29 (0.08) |
| spruce | $\hat{d}^2_{ind}$ | 0.03 (0.11) | - | 0.31 (0.10) | -0.00 (0.08) |
| | $\hat{H}^2_{ind}$ | 0.08 (0.11) | - | 0.59 (0.10) | 0.28 (0.08) |

Type-B genetic correlations were weaker for most growth traits, indicating important genotype-by-environment interactions for this category of traits; while stronger genetic correlations were recorded for wood quality traits, in line with the generally higher heritability estimates obtained for this category of traits. Also, growth traits showed significant dominance effects, whereas these effects were usually less pronounced for wood quality traits. These findings underscore the distinct genetic architectures between growth and wood quality traits, and they emphasize the role of site-specific interactions in shaping trait expression in a heterogenous manner.

For the reporting of the overall trends, we focus here on the across-site heritability estimates. The repetitive sampling of SNP subsets and their use in GS models (GBLUP-AD) allowed describing trends in heritability estimates, model predictive ability, and prediction accuracy as a function of number of markers used, and comparing with the estimates obtained from models using all available markers and those with pedigree information only (ABLUP-AD,) for the various growth and wood quality traits in the three studied species.

As a general trend, narrow-sense heritability and the dominance ratio of growth and wood quality traits appeared to rapidly increase from between 500 and 4000 markers (Figs. 1 to 3). However, starting from about 4000 SNPs, the rate of change in narrow-sense heritability and dominance estimates became substantially lower compared to the sharp rise in trend observed before this threshold. Heritability reached a plateau after 8000 SNPs for black spruce and white spruce, and at a slightly higher level in Norway spruce. It is also notable that the reduction of standard deviation of estimates after 4000 SNPs was also much limited. Thus, the marginal improvement of heritability estimates between 4000 and 8000 SNPs does not seem to justify efforts to double marker density unless this is useful to improve modeling accuracy significantly (see Discussion). A few exceptions were noted. For instance, a slight increase of broad-sense heritability estimates was detected beyond 10,000 SNPs for acoustic velocity in white spruce (Fig.2), which was linked to a small increase in additive genetic variance estimates, although this increase remained marginal in the range of a few percent improvement. For Norway spruce (Fig.3), the rise of heritability estimates beyond a few thousand markers was also minimal. From 25,000 SNPs onwards, the heritability estimates remained stable up to 100,000 SNPs. In this Norway spruce dataset, heritability estimates for tree height were consistently very low, regardless of the size of SNP subsets tested.

**Fig. 1.**
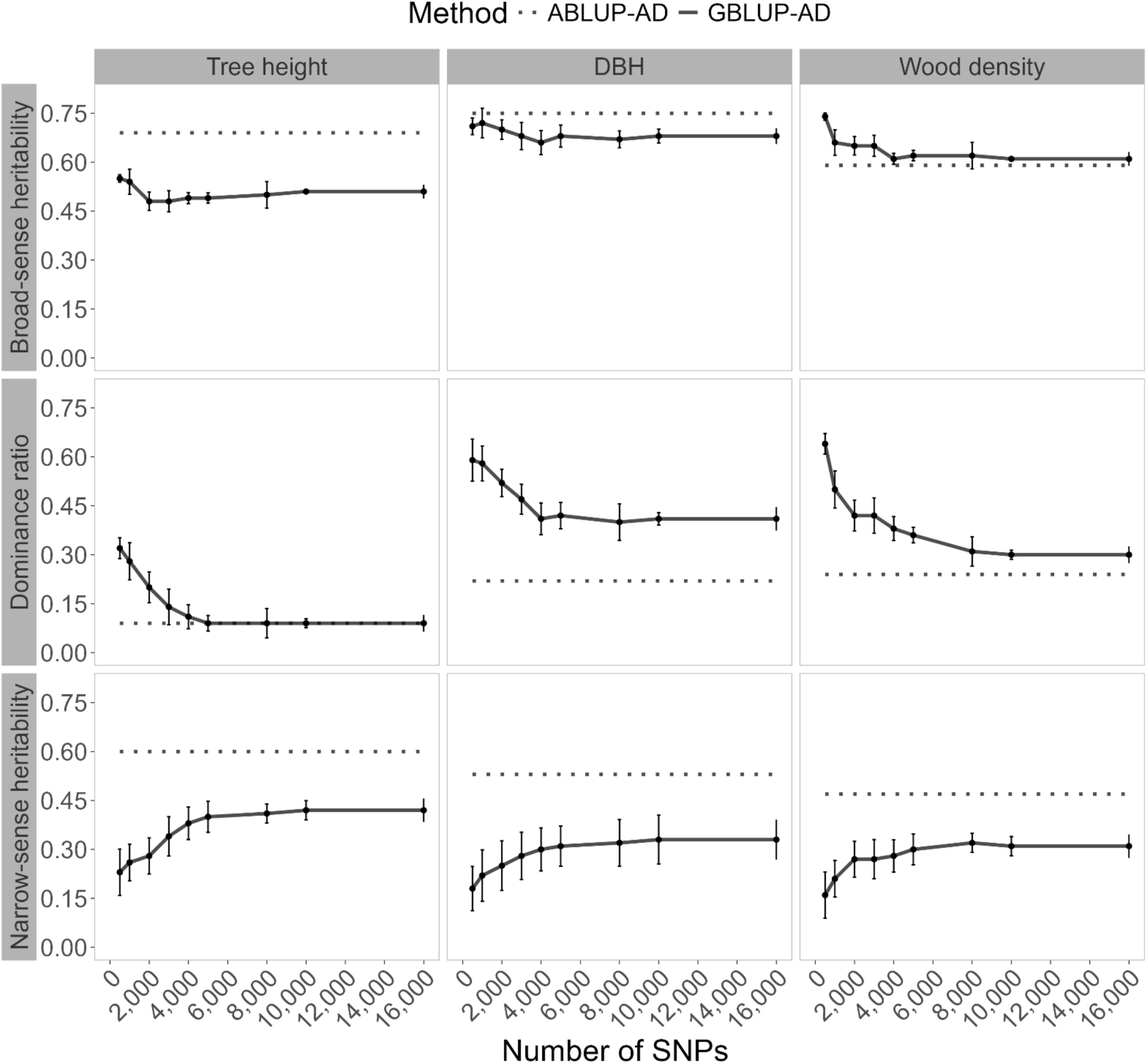
Black spruce heritability trends in function of the number of markers used in GS modeling of tree height, DBH, and wood density. Broad-sense heritability and its components, the narrow-sense heritability and dominance ratio are shown in horizontal panels. Solid lines present the mean estimates of SNP resampling of GBLUP-AD models, including the standard deviations. The dashed lines represent estimates from the corresponding pedigree-based models (ABLUP-AD). DBH is tree diameter at breast height.

**Fig. 2.**
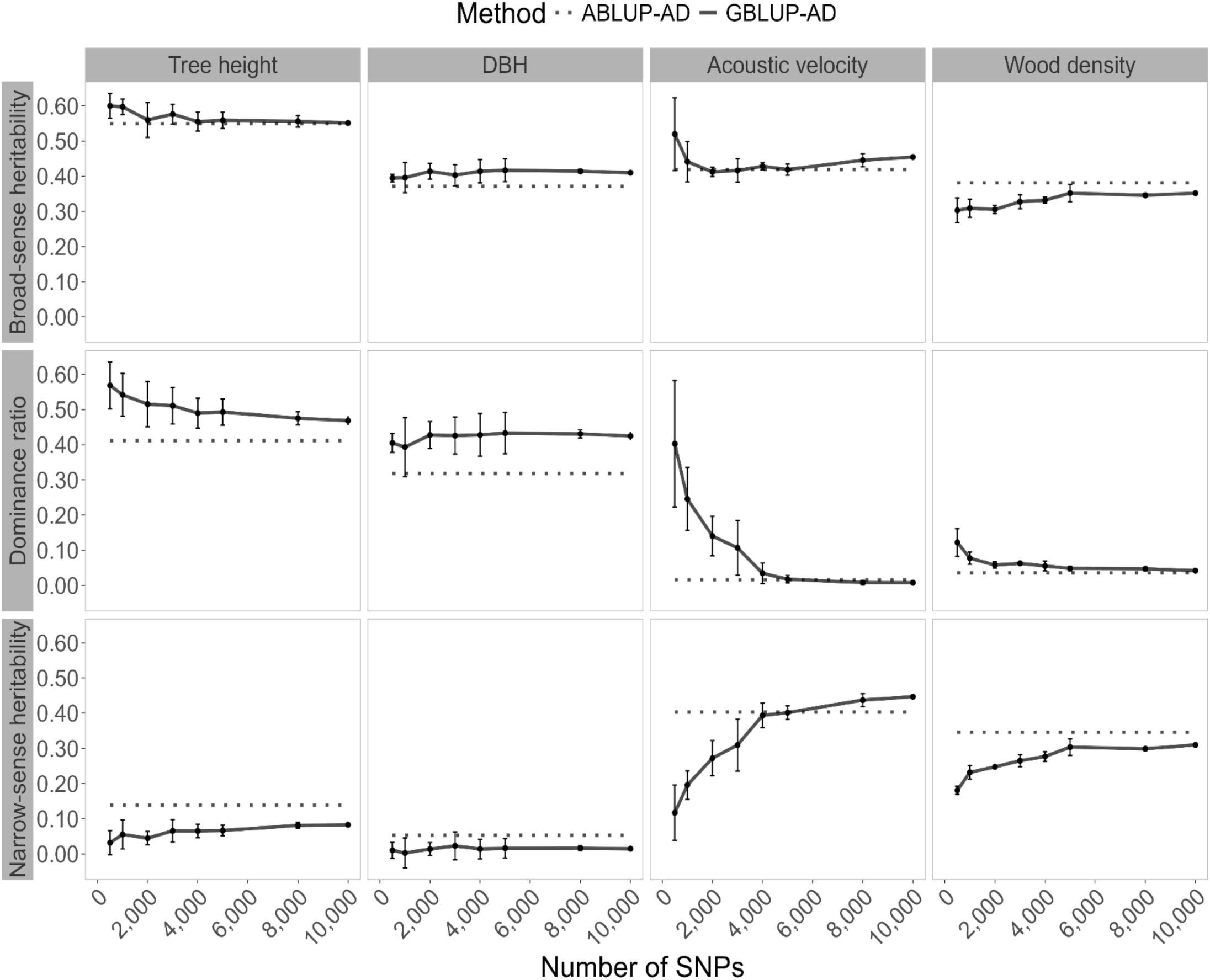
White spruce heritability trends in function of the number of markers used in GS modeling of tree height, DBH, acoustic velocity, and wood density. Broad-sense heritability and its components, the narrow-sense heritability and dominance ratio are shown in horizontal panels. Solid lines present the mean estimates of SNP resampling of GBLUP-AD models, including the standard deviations. The dashed lines represent estimates from the corresponding pedigree-based models (ABLUP-AD). DBH is tree diameter at breast height.

**Fig. 3.**
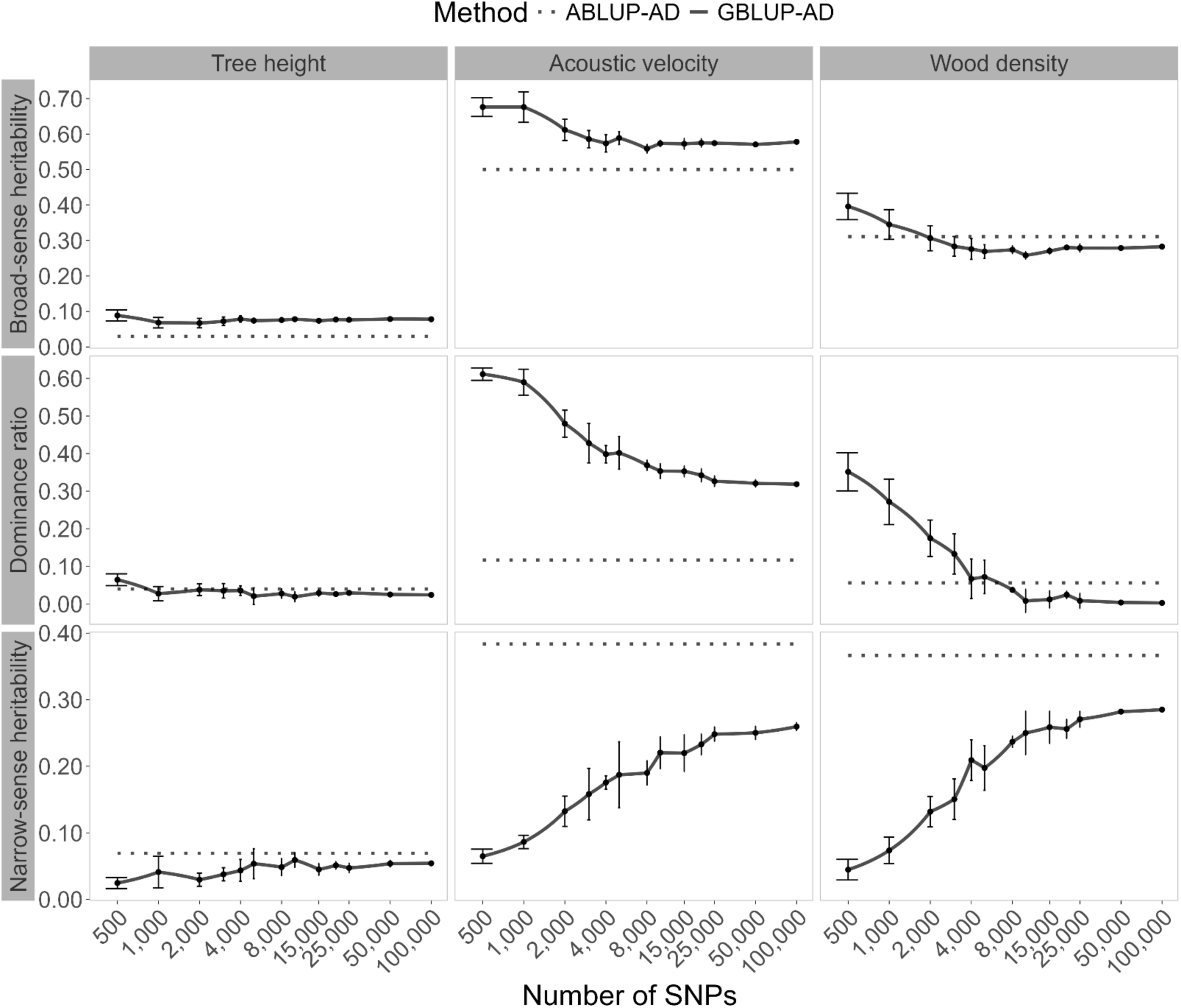
Norway spruce heritability trends in function of the number of markers used in GS modeling of tree height, acoustic velocity, and wood density. Broad-sense heritability and its components, the narrow-sense heritability and dominance ratio are shown in horizontal panels. Solid lines present the mean estimates of SNP resampling of GBLUP-AD models, including the standard deviations. For a better readability, a logarithmic scale is used in this figure for the number of SNPs in marker subsets. The dashed lines represent estimates from the corresponding pedigree-based models (ABLUP-AD). DBH is tree diameter at breast height.

In contrast, broad-sense heritability was less variable at low marker density, compared with the respective dominance component and narrow-sense heritability estimates in all three species. The estimated dominance ratio using GBLUP-AD was initially quite high when low marker density was considered in GS models, but tended to decrease as marker density increased, ultimately converging toward the dominance ratio observed in ABLUP-AD models. Models based on low marker density were apparently less performant in separating additive from dominance effects. Levelling out of estimates from GBLUP-AD models at higher numbers of SNPs went along with reduced standard deviations and slight rise of parameter estimates after 4000 SNPs, and virtually no change after 8000 SNPs.

The trends of mean predictive ability of breeding values (*PA_BV_*) and of genetic values (*PA_GV_*) as a function of the number of SNPs used in GS models are illustrated in Fig. 4 for black spruce, and for white spruce and Norway spruce, in Fig. 5 and Fig. 6, respectively. Across all species, an increase in predictive ability was observed as marker density increased and this, for both predictive ability of breeding values and genetic values. Mean estimates tended to increase to a maximum between 4000 and 8000 SNPs, and remain stable thereafter. Similarly, as for the trends in heritability estimates in Norway spruce, a minimal increase of predictive ability was observed beyond 4000 SNPs, reaching a plateau at around 25,000 SNPs. The trends observed for predictive ability were very similar to the trends observed for the prediction accuracy of breeding values (*PACC*_*BV*_) and of genetic values (*PACC*_*GV*_) (See Supplementary, Fig. S1-S3). In the interest to describe trends in function of marker density used in GS models, we focus primarily on the trends in predictive ability estimates below.

**Fig. 4.**
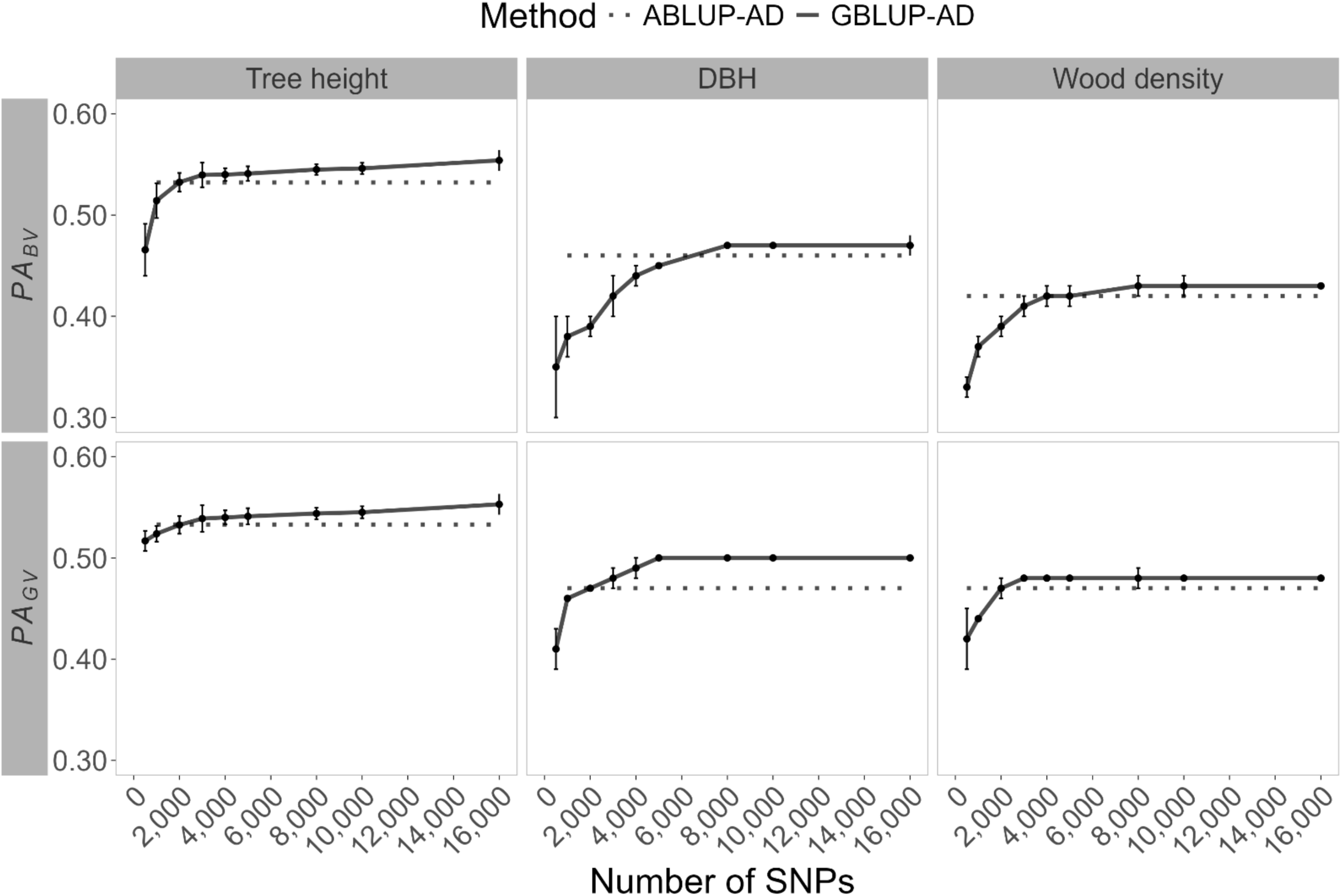
Trends in average predictive ability of breeding values (***PA***_***BV***_) and genetic values (***PA***_***GV***_) in black spruce GS models (GBLUP-AD) considering varying subsets of SNPs. For the three phenotypic traits, the means and standard deviations are shown for ten-fold cross validation of ten times resampled marker subsets. Horizontal dashed lines represent the corresponding pedigree-based models (ABLUP-AD), independent of SNP number. DBH is tree diameter at breast height.

**Fig. 5.**
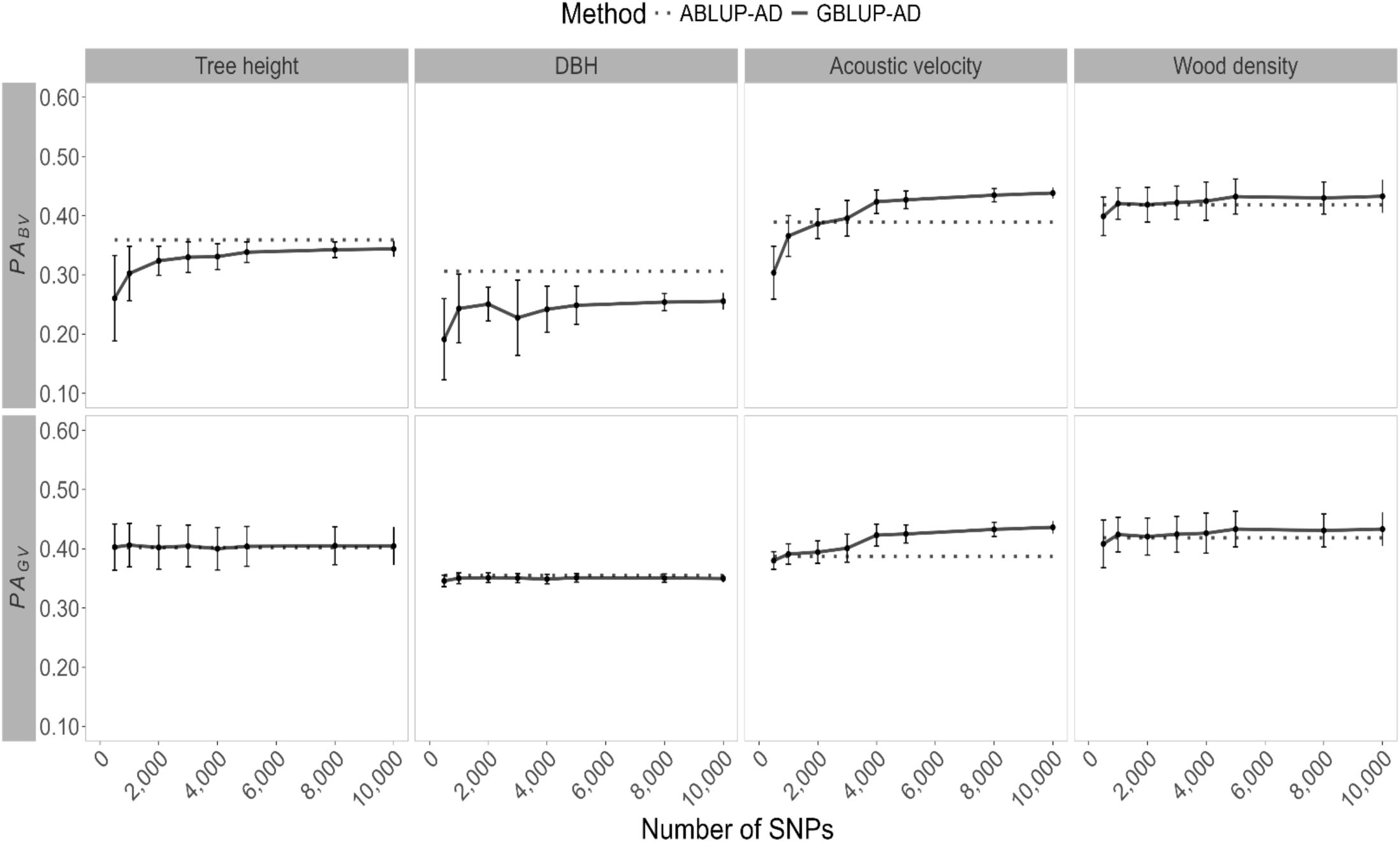
Trends of average predictive ability of breeding values (***PA***_***BV***_) and genetic values (***PA***_***GV***_) in white spruce GS models (GBLUP-AD) considering varying subsets of SNPs. For the four phenotypic traits, the means and standard deviations are shown for ten-fold cross validation of ten times resampled marker subsets. Horizontal dashed lines represent the corresponding pedigree-based models (ABLUP-AD), independent of SNP number. DBH is tree diameter at breast height.

**Fig. 6.**
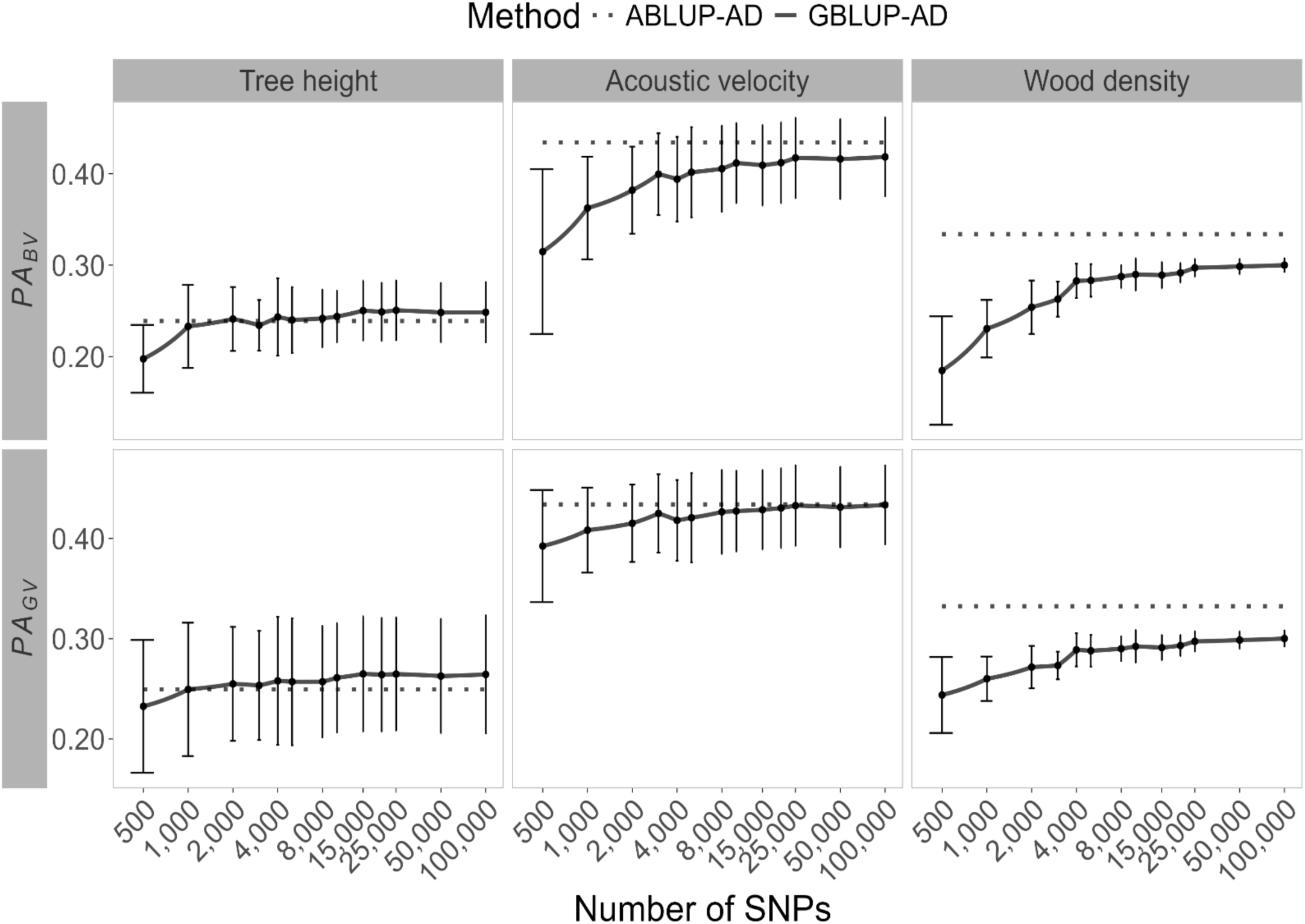
Trends in average predictive ability of breeding values (***PA***_***BV***_) and genetic values (***PA***_***GV***_) in Norway spruce GS models (GBLUP-AD) considering varying numbers of SNPs. For the three phenotypic traits, the means and standard deviation are shown for ten-fold cross validation of ten times resampled marker subsets. Horizontal dashed lines represent the corresponding pedigree-based models (ABLUP-AD), independent of the number of SNPs. The number of SNPs is presented on a logarithmic scale to facilitate readability and interpretability of results obtained for smaller marker subsets.

In a very few cases, a slight decrease of predictive ability was observed around 2000 or 4000 SNPs despite the general increasing trend. This was so for DBH in black spruce and white spruce, although these estimates were characterized by relatively large standard deviations overlapping with the mean estimates of predictive ability obtained for the neighbouring smaller and larger SNP densities tested.

In general, predictive ability of GS models tended to be higher or very close to pedigree-based predictive ability estimates. Wood density in Norway spruce presented thereby an exception. as well as growth traits in white spruce, where the pedigree-based predictive ability was greater than that obtained with the genomic models (GBLUP-AD). Similarly, as for trends in heritability estimates, predictive ability of genetic values, i.e., including additive and dominance effects (*PA_GV_*), tended to vary less than their breeding value counterparts that only considered additive effects. This trend again hints towards the insufficiency of GS models relying on low marker densities to separate well additive and dominance effects, and it underlines the need to consider non-additive effects in GS modelling whenever possible.

## 4 Discussion

During the last two decades, the use of EST and the progress in RNA- and DNA-based sequencing methods along with the development of automated computing pipelines to detect and to filter single nucleotide polymorphisms (SNPs) from gene sequences, has allowed the creation and functional annotation of significant marker resources for spruces (Pavy et al. 2006, 2016, 2025; Chen et al. 2012; Nystedt et al. 2013; Azaiez et al. 2018; Bernhardsson et al. 2019; Gagalova et al. 2022; Lo et al. 2024). Those resources have permitted the construction of genetic maps (Pelgas et al. 2006, 2011; Lind et al. 2014; Pavy et al. 2012, 2017; Bernhardsson et al. 2019; Gagalova et al. 2022) as well as the creation of new genotyping tools for breeding applications in white spruce (Pavy et al. 2008, 2013; Lenz et al. 2020a), black spruce (Pavy et al. 2016) and Norway spruce (Lenz et al. 2020a; Bernhardsson et al. 2021). Still, the use of many thousand markers for the construction of SNP genotyping arrays and the subsequent genotyping of thousands of trees is associated to significant costs. Consequently, it is perceived by some tree breeders as a financial burden to the large-scale application of GS in tree breeding programs (Grattapaglia 2022), perhaps ignoring the large-scale benefits pf GS per unit of time to accelerate the deployment of improved varieties in reforestation programs and improved plantations (Chamberland et al. 2020), which are downstream benefits often not considered in the funding of tree breeding programs.

Based on major datasets, the results of the present study appear as a significant step forward to inform on the number of markers needed for most exact estimation of genetic parameters such as heritability and the precision of GS models for the prediction of breeding values, at least in conifer species with very large giga-genomes such as for spruces (see Beaulieu et al. [2024] Table 1). Furthermore, a well-defined selection of marker panels appears essential for operational decisions regarding genotyping for GS applications, as breeding program managers may have to balance accuracy of parameter estimates and associated costs for genotyping. Although metadata analysis and simulation studies such as Grattapaglia and Resende (2011) and Beaulieu et al. (2022, 2004) have highlighted key elements impacting the precision of GS models and genetic parameter estimates, comprehensive empirical studies that demonstrate directly these effects at the early-stage of tree breeding programs are still scarce.

We focused our analyses on broad- and narrow-sense heritability to assess the models’ capacity to estimate and separate additive and non-additive effects, as well as their effects on predictive ability (*PA*) and prediction accuracy (*PACC*) to evaluate the ability of GS models to accurately predict breeding and genetic values using of variety of marker densities. For all three spruce species, general estimates based on pedigree information, or based on all available markers were in the same range as those reported in earlier homologous studies by Lenz et al. (2017) for black spruce, Beaulieu et al. (2014b) for white spruce, and (Chen et al. 2018) for Norway spruce. Some mostly minor differences were related to the fact that different algorithms and models were used. The present study relied on a linear mixed model framework (GBLUP) as it represents a straightforward way to consider heterogenous error and genetic variances across different environments, and to include dominance effects in these controlled-cross populations made of related full-sib families. For black spruce and white spruce, many more markers were available than in previous studies, which more than doubled the number of markers used compared with the initial publications (Lenz et al. 2017 and Beaulieu et al. 2014b, respectively), thus allowing to reach plateaus in estimates for most studied parameters derived from GS, as for Norway spruce.

### 4.1 Inffuence of marker density on genetic parameter estimates

The subsampling approach used herein to randomly create subsets of markers differing in numbers allowed us to obtain the trends in heritability, *PA* and *PACC* estimates with increasing marker density. The general trends were largely mirrored across the three species despite some differences in experimental setups and age of trait measurements among species. Heritability and predictive ability increased significantly among subsets varying from 500 to around 4000 SNPs, and low improvement or rather stagnation of estimates was observed beyond. Additionally, standard deviation of estimates decreased as the number of SNPs increased, reinforcing the notion that reliable heritability estimates are highly dependent on the quality and size of the SNP dataset.

It is difficult to conclude about the reason for the slightly later culmination of heritability and accuracy estimates in Norway spruce around 25,000 SNPs, as this could be related to the different experimental design. Given the more intensive sampling of SNPs per gene locus to build the Norway spruce SNP array (Berhardsson et al. 2018) than for the other two species, on average, genes with many SNPs arrayed were more likely to be retained in the small subsets of SNPs that those represented by few SNPs, sometimes in a redundant manner, thus reducing marker density in terms of distinct sampled loci compared to black spruce or white spruce, with less redundant SNPs per genetic locus on the SNP array. But more importantly, the Norway spruce dataset contained more parents and families (thus higher status number) but a smaller number of trees tested per family, than in the black spruce or white spruce datasets (Supplementary Table S4). This may have lead to less precise estimation of genetic variance at lower values of marker density tested.

In line with this, a recent meta-analysis (Beaulieu et al. 2024) and several other simulation (Grattapaglia and Resende 2011) and empirical studies reported a negative relationship between status number and predictive ability such that, as a general trend, breeding values could be more precisely estimated in populations of smaller effective size for about the same range of number of trees considered. They also found no relationship between heritability estimates and parameters such as the number of markers or the size of training set in the range of values available from the literature (Beaulieu et al. 2024). This also underlines that heritability is a site- and population-specific measure, meaning that broad trends can be more difficult to detect when analysing data from varied population structures and study designs. These findings suggest caution when extrapolating results from the present study to very different genetic structures of breeding populations, such as for breeding programs based on half-sib families from natural populations, which usually harbor much larger census and effective sizes than populations of related full-sib families, such as analysed here for these three spruce species (see also Beaulieu et al. [2014a] vs. [2014b] for comparison of GS accuracy in context of half-sib versus full-sib population structure in same species with similar GS modeling).

Earlier studies had already indicated that low marker density, in the range of a few thousand markers, can lead to reasonable prediction accuracy of genomic-estimated breeding values in the context of related full-sib families and small effective sizes (Beaulieu et al. 2014b; Lenz et al. 2017). The resampling approach used in this study also allows generalizing such observations made on Norway spruce by Chen et al. (2018). The findings that available total marker panels were not necessary to achieve precise estimates of genetic parameters and GS accuracy underscore the diminishing returns of adding more SNPs beyond a certain threshold, emphasizing the importance of an optimal balance considering dataset size, population structure and GS accuracy. Similar results were reported in *Eucalyptus benthamii* Maiden & Cambage, a broadleaves tree species, where Estopa et al. (2023) showed that a relatively small number of SNPs (less than 10,000 for most traits) was sufficient to obtain accurate parameter estimates for growth and wood quality traits. Our findings are also consistent with reports on the influence of marker density on genomic prediction accuracy in plant and animal breeding, such as in alfafa (Wang et al. 2024), maize (Guo et al. 2020), wheat (Norman et al. 2018), cattle and pigs (Zhang et al. 2019). Those studies showed that increasing the number of markers led to higher prediction accuracy, but only up to certain thresholds beyond which additional markers provided limited enhancement. This pattern mirrors observations from trend analysis of results gathered from many GS studies in forest tree species where a significant relationship was not observed between marker density and GS accuracy (Beaulieu et al. 2024). Altogether, these studies suggest the relevance of identifying an optimal marker density that balances genotyping costs with genetic parameter and prediction accuracy, particularly when considering application of GS in often resource-limited tree breeding programs.

As reported by Estopa et al. (2023) for *E. benthamii*, the earlier culmination and plateau of values reached for *PA* in relation to an increasing number of markers, compared to that for heritability estimates, was not observed as a clear trend in the present study on three spruce species. Measures of GS model precision, such as *PA,* were rather following closely trends in heritability estimates, which necessitated a marginally lower marker density for traits with low heritability to attain plateaus of values. Given that quantitative traits with weak or weaker genetic control have a smaller proportion of their phenotypic variation explained by genetic effects, these traits might necessitate less marker density for their heritability to be traced adequately, and that most QTLs bear small gene effects such as those in relation to adaptive traits (Pelgas et al. 2011). Further studies are needed on comparative genomic architecture between quantitative traits with low versus high heritability in tree species.

### 4.2 Higher marker density helped disentangle additive from non-additive genetic effects

Pedigree-based models tended to result in slightly higher heritability estimates than those derived from GS models, which was especially so for narrow-sense heritability estimates. For example, for black spruce growth traits, narrow-sense heritability estimates were higher in pedigree-based models compared to their marker-based counterparts. At the same time, those higher estimates for pedigree-based models did not lead to higher or significantly higher *PA_BV_* values. Similar findings were reported by Chen et al. (2018) for wood quality traits in Norway spruce, where the pedigree-based ABLUP models also produced higher heritability estimates. This result may be explained by the fact that the pedigree-based models can overestimate additive variance, and consequently these heritability estimates are not relying on actual genetic relationships (Beaulieu et al. 2022). Comparable results were observed in *E. benthamii* (Estopa et al. 2023), in black spruce (Lenz et al. 2017) and in a meta-analyses of GS applied in various tree species (Beaulieu et al. 2022, 2024). GS models are thus expected to provide more accurate estimates of variance components and heritability compared to pedigree-based models, especially when plateaus of values have been reached. Indeed, pedigree-based methods overlook genetic relationships beyond those recorded in the registered pedigree and thus, rely on expected rather than actual genetic relationships. Furthermore, these models assume homogeneity in relationships among the same types of parents, which means that genetic covariance components are estimated primarily from interfamilial variation. This approach has limitations because Mendelian sampling deviations cannot be distinguished from non-genetic residual errors (Ødegård and Meuwissen 2012). Moreover, if errors exist in the registered pedigree, they cannot be corrected, contrary to GBLUP for instance (Godbout et al. 2017; Li et al. 2019).

Experimental studies have shown that incorporating dominance effects into genomic models reduces the estimated additive genetic variance and narrow-sense heritability (De Almeida Filho et al. 2019; Paludeto et al. 2021; Nadeau et al. 2023). Indeed, dominance effects represent a portion of genetic variance that appears to be less well-captured by pedigree-based models than marker-based models. By better capturing non-additive genetic components, GS models provide more accurate estimates of additive variance, often leading to lower heritability estimates compared to pedigree-based models (Beaulieu et al. 2022). This improvement in accuracy highlights the importance of accounting for non-additive effects in genetic analysis, whenever possible.

Furthermore, almost all GS models lead to the highest *PA_GV_* values when dominance deviation was included in the prediction of genetic values. This trend is also supported by other studies in white spruce (Nadeau et al. 2023). For spruce and tree breeders aiming to estimate genetic parameters of their breeding populations, that is, including both additive and dominance effects, our results indicate that this can also be done at relatively low marker densities, between 4000 SNPs and 8000 SNPs. At the lowest end of this range however, additive and dominance variances do not appear to be well disentangled, which may lead to less precise estimates of genetic values and suboptimal selection of breeding and reproduction material. Nevertheless, *PA_GV_* values were already high at quite low marker density, indicating that genetic values can already be estimated with sufficient accuracy with quite low marker densities. But this is of rather restricted use for several conifer breeding programs, given that only few programs apply clonal reproduction by rooted cuttings or somatic embryogenesis (SE) for the production of improved varieties, where non-additive genetic effects can indeed be considered and exploited for additional genetic gains (Park et al. 2016).

### 4.3. Genomic distribution of markers and genome architecture

Our results indicated that only moderate marker densities are needed to attain high precision of genetic parameter estimates and accuracy of GS models for an array of functional traits, as applied to spruce breeding populations of relatively small status number typical of advanced-breeding populations. One must remember that such breeding populations are characterized by high LD, contrary to outcrossing natural populations of forest trees usually with large effective population sizes Bouillé and Bousquet 2005) where LD decays rapidly, often well within gene limits (Neale and Savolainen; Pavy et al. 2012b). For instance, in genome-wide association studies (GWAS) such as the detection of genotype-phenotype associations (GPA) or genotype-environmental associations (GEA), where significant associations are mostly determined by short-range LD between markers and QTLs, unstructured populations or structured populations stripped of most of their genetic structure must be usually used in GWAS to avoid artifactual results (e.g. Hornoy et al. 2015; Depardieu et al. 2021). In such cases, very large numbers of markers and genetic loci need to be screened to identify significant associations, given the rapid decay of LD in these populations. In breeding populations where GS is usually applied, especially those made of related families and of reduced status number as in advanced-breeding generations, they are more highly structured with high LD, making GS most accurate while relying largely on relatedness instead of short-range LD between markers and QTLs (Habier et al. 2007; Zapata-Valenzuela et al. 2012; Beaulieu et al. 2014b; Lenz et al. 2017; Grattapaglia et al. 2022).

While GS model accuracy largely depended on the number of markers used in the context of capturing well relatedness, it is also likely that the number of distinct genetic loci considered in SNP array development or any other genotyping method, and their distribution on the genome, also plays a significant role in the efficiency of the marker density used to capture relatedness with more reduced numbers of markers. Indeed, the white spruce and black spruce SNP arrays used herein relied on maximizing as much as possible the number of distinct gene loci on the array so to improve genome coverage for a same number of markers, thus with a reduced SNP redundancy of less than two SNPs per gene locus, on average. As a test control, many more SNPs per gene locus were arrayed and tested for Norway spruce, without clear improvement of GS accuracy well beyond the number of gene clusters included in the SNP array (around 20,000). This trend indicates that SNPs within same genes were generally in high LD in such structured breeding populations of small status number, with on average diminishing GS returns of the individual informative value of each SNP when being redundant in same gene loci.

A quite even distribution of markers onto the genome also appears important, which is often an information not available for many tree species. For instance, in the present study, loci sampled in each species for SNP array design were distributed exome-wide, with previous studies indicating that the exome covers quite uniformly the 12 linkage groups or chromosomes of spruce sp. (Pavy et al. 2008, 2012, 2017; Lind et al. 2014; Gagalova et al. 2022). This is likely to be the case for most tree species. It was further shown by Lenz et al. (2017) that when limiting GS modeling to gene SNPs from one chromosome at a time, model accuracy was below that obtained when considering the same number of SNPs but sampled from all 12 spruce chromosomes. It was suggested that relatedness could not be entirely captured when limiting marker sampling to one chromosome, or more likely, that short-range marker-QTL LD was significant but could not be entirely captured when marker distribution was skewed toward only a portion of the genome. Future efforts to design SNP genotyping arrays or to orientate genotype-by-sequencing (GbS) strategies should thus consider not only the total number of SNPs required to optimize genome coverage and GS model performance, but also the distribution and number of distinct genetic loci sampled. A thoughtful selection of markers representative of a large array of genetic loci and thus, with a genome-wide or exome-wide distribution should help improve the balance between genome coverage in markers, GS precision and genotyping costs so to optimize and make cost-effective the application of GS in a given breeding program.

It was further informed by Beaulieu et al. (2024) that irrespective of tree species genome size in base pairs, which may vary by a factor of more than 50 times among tree species (from about 0.5 Gb to above 20 Gb), that the genome size in centimorgans (cM), thus based on recombination frequency, was of the same magnitude for most tree species so far targeted by GS. This suggests, for similar breeding population structures, that the marker density for most tree species should be in the same range, in order to reach plateaus of genetic parameter estimates and GS model accuracy. Thus, the optimal levels marker density identified in the present study, in terms of number of markers per cM of around two to four for spruce genomes of about 2000 cM (Beaulieu et al. 2024), should be applicable to most conifer and broadleaves tree species in a similar range of genome size in cM. Such a number of markers and marker density is also well in agreement with early simulation results of Grattapaglia and Resende (2011) and the empirical results of Estopa et al. (2023) in *E. benthamii*, which is characterized by a much smaller physical genome that that of spruces or conifers in general, but of same magnitude of genome size in cM (using *E. nitens* as a baseline reference for *Eucalyptus* sp., see Beaulieu et al. [2024] Table 1).

Preselection of markers has also been reported to enhance GS model performance (Lenz et al. 2017; Chen et al. 2018; Thumma et al. 2022), which could thus serve to enhance model accuracy when only the manufacture of small SNP arrays could be afforded, or to reduce the number of well-distributed markers needed to reach plateaus of model accuracy, given the type of breeding population targeted. Depending on the selection method, this could possibly shift the usual reliance of GS model accuracy on relatedness so to enhance the detection of marker-QTL LD.

High LD has also been suggested to make GS model predictions more accurate and sustainable, not only across related populations, but also across generations (Habier et al. 2010, 2013). In the context of long-term breeding programs, such as those in animal genetics or for forest trees, deeper pedigrees and accumulated recombination events could erode linkage between markers and QTLs, though this would less affect long-range LD underlying relatedness. Indeed, preliminary demonstrations in conifers are very encouraging when relatedness is maintained across generations (Bartholomé et al. 2016). This question may require more studies using haplotype-based approaches to efficiently capture and track their structures across generations.

### 4.4. Breeding applications

Most gain per unit of time and economic benefits in tree breeding could be accomplished by combining GS with multiclonal selection and deployment (Park et al. 2016; Chamberland et al. 2020; McLean et al. 2023) whenever possible, which is particularly amenable in species prone to vegetative propagation such as spruces. With forward selection, best clones can be selected within families for propagation if good separation of additive from non-additive effects is possible, as shown in this study. Currently, tree breeding programs rely mostly on additive genetic effects, though the situation is changing with the evolution towards more advanced-breeding generations in many tree breeding programs (Mullin et al. 2011; Li and Dungey 2018; Lenz et al. 2020a,b; Grattapaglia et al. 2022).

Given that once manufactured, SNP genotyping arrays are re-usable for many years and deliver stable genotyping results (Bousquet et al. 2021), or that they could be designed efficiently for the needs of several closely-related species (Silva-Junior et al. 2015; Gérardi et al. 2025), the popular belief is still that they remain too costly for tree breeding programs that often operate with limited budgets. However, one must also realize that the genotyping costs, including those for the manufacture of such SNP genotyping arrays, have decreased significantly since the introduction of GS in breeding (Gorjanc et al. 2015; Bhandari et al. 2019). Carefully designed lower-density SNP arrays should also provide a more affordable solution to tree breeders, with final genotyping cost per sample decreasing with increase in the number of samples expected to be genotyped and thus, planning ahead for multi-year needs by the tree breeders.

The present study revealed that a well-distributed number of markers of around 4000 to 8000 SNPs for a genome size of around 2000 cM would (corresponding to 2 to 4 SNPs/cM) resulted in nearly-maximum heritability estimates and GS model precision for both growth and wood quality traits. Our results indicate that additional genotyping costs aiming at further improving GS estimates beyond such a marker density may not be necessary, as additional gains will most likely be marginal even by doubling or tripling marker density. Consequently, such marker density for advanced-breeding populations of relatively small status number offers a fair trade-off between genotyping costs and the precision of genetic parameters and GS model accuracy obtained, making further investment in genotyping less justifiable. Beyond such a level of marker density, it is also more likely that improving size of training set in GS modeling (e.g. Nadeau et al. 2023) will be more productive than increasing the number of markers by a large factor. This is because of the presence of high LD in advanced-breeding populations, which usually relies on controlled crosses or polycross (e.g. Lenz et al. 2020b) of a quite limited number of selected parents, and the necessity of sampling well within families to maximize modeling accuracy, whether by conventional pedigree methods or by GS.

Thus, the remaining main obstacle to fully integrating GS into existing tree breeding programs does not appear technical anymore, but mostly of strategic and financial nature. As emphasized by Grattapaglia et al. (2022) and several others, identifying well the objectives and understanding the logistic and financial dimensions of implementing GS in a given tree breeding programs appears to be crucial for its adoption and success. A careful balance between genotyping costs and GS model accuracy should thus contribute to ensuring the long-term adoption of GS in tree breeding and hastening breeding cycles in the context of rapid environmental change (Bousquet et al. 2021; Laverdière et al. 2022). As such, unconvinced tree breeders should have a second look at the costs and benefits of implementing GS in their breeding programs, especially with a vision of mid- to long term needs and applications.

## 5 Conclusions

Our study tested varying marker density levels and their effects on the accuracy of GS models for traits of different genetic architecture in three boreal conifers widely reforested in the northern hemisphere. The results bear important implications for the operational deployment of GS in tree breeding programs beyond those of black spruce, white spruce, and Norway spruce. Despite some differences in experimental setups among the cases analysed herein, similar plateaus in heritability estimates and predictive ability of the GS models were observed between 4000 and 8000 SNPs, which were largely representing distinct gene loci well spread on the genome. This trend indicates that this range of relatively low marker density may offer a practical balance between GS accuracy and cost, although albeit additional gains may still be achievable with larger numbers of markers in populations with much larger status numbers, but usually with diminishing rewards as genetic structure is reduced and status number increases. Given the high cost of manufacturing and using very large SNP arrays, particularly in tree breeding contexts where budgets are often much constrained, smaller exome-wide SNP panels designed to capture maximum genetic information per marker used should provide a cost-effective alternative, especially for applications in breeding populations of small status number where GS accuracy is mostly relying on relatedness. Thus, the need for surveying very large number of SNPs, as usually needed for GWAS applications in unstructured or natural populations of conifers of large effective population size and low levels of LD, should certainly not be taken as a yardstick for GS applications, given the high recombination rates and rapid LD decay across many generations in these populations (Bousquet et al. 2021). This is contrary to advanced-breeding populations with usually much smaller status number and more relatedness, where GS should be most effectively applied, thus necessitating much less intense genome coverage.

Our results also underscore the importance of considering the diversity of sampled genetic loci and their genomic distribution when designing genotyping arrays or for GbS strategy. Because of SNPs generally in high LD within genetic loci among related individuals, these results demonstrate the relevance of reducing SNP redundancy within loci and relying on relatively low numbers but well-distributed markers on the genome for predictive performance within a single generation of GS. Operational implementation of GS should thus aim for cost efficiency by maximising marker distribution on the genome for a given allocation of SNPs in genotyping strategies. As genotyping technologies have become more accessible and reliable, strategic planning and size of financial investment rather than technical feasibility are now representing the key factors determining the feasibility and successful routine integration of GS into forest tree breeding programs.

## Supporting information

Supplementary Materiel

## Statements and Declarations

The authors have no competing interests to declare that are relevant to the content of this article.

## Funding

This research was supported by multiple funding agencies across Canada and Sweden. The Ministère des Ressources naturelles et des forêts of Québec supported the maintenance of black spruce and white spruce field tests and sampling, while the Canadian Forest Service and the Natural Resources Canada Genomics RCD Initiative contributed resources to data management, computational resources, and analytical support. SNP genotyping for black spruce and white spruce were funded by Genome Canada and Genome Quebec through grants supporting the FastTRAC II genomic project led by J. Bousquet and P. Lenz, and additional support was received through the Canada Research Chair in Forest Genomics (Univ. Laval). For the Norway spruce component of the study, financial support was provided to H.X. Wu by the Swedish Research Council for Environment, Agricultural Sciences and Spatial Planning (Formas; grant 230–2014-427) and the Swedish Foundation for Strategic Research (SSF; grant RBP14–0040). The Swedish nor the Canadian funders had any role in study design, data collection, analysis, or publication strategy.

## Data Availability Statement

In order to comply with Intellectual Property Policies (IPP) of participating governmental and private organizations in this work, the supporting phenotyping and genotyping data is not deposited into the public domain. However, data will be shared upon motivated request to the corresponding author after consulting with participating governmental and private organizations involved in this work.

## Acknowledgements

We thank all collaborators and institutions involved in the establishment, maintenance, and data collection of the various spruce trials used herein. For the black spruce component, we are grateful to M. Villeneuve (formerly at the Direction de la recherche forestière of Ministère des Ressources naturelles et des forêts of Québec, DRF-MRNF) for initiating the black spruce biparental test over 25 years ago, and to G. Gagnon, G. Numainville, and several others from DRF-MRNF for their work on controlled crosses, field sampling and phenotyping. For the white spruce component, we acknowledge the contributions of G. Daoust, R. Paquet, D. Plourde, the late S. Légaré (Natural Resources Canada), A. Rainville, G. Gagnon (DRF-MRNF) and their teams for establishing and maintaining the trials, as well as E. Dussault (Natural Resources Canada) and collaborators for phenotypic and genomic data collection. We also thank S. Gérardi and N. Pavy (Canada Research Chair in Forest Genomics, Univ. Laval) for developing the SNP resources and their roles in designing the black spruce and white spruce genotyping SNP arrays. We also acknowledge F. Bacot and F. Robidoux and their team at the Genome Quebec Innovation Centre (Montreal, Canada) for their support for SNP array manufactures and conducting the Infinium genotyping assays. For the Norway spruce component, genetic experiments (test plantations and clonal archives) were established and maintained by the Forestry Research Institute of Sweden (Skogforsk) for breeding selections and research purposes. Computations were performed using resources provided by the Swedish National Infrastructure for Computing (SNIC) at UPPMAX and HPC2N. We also thank Dr. Zhiqiang Chen, Junjie Zhang, Tianyi Liu, Xinyu Chen, Linghua Zhou, and Anders Fries (Umeå Plant Science Centre) for their contributions to design of the Norway spruce SNP genotyping chip, DNA extraction and genotyping of Norway spruce samples, field assistance, trial maintenance and quantitative data.

## Authors’ Contributions

A.S. assisted in designing the study, curated and analysed the data, conducted analyses, and prepared the original draft of the manuscript. J.Bo., P.L and JBe. contributed to conceptualization, study design, investigation, validation, supervision, funding support, and manuscript reviews. J.-P.L. and S.N. contributed to data analysis and manuscript review. F.G. handled the new raw genotyping data for black spruce and white spruce and validation of their quality, also contributing to manuscript review. F.O. contributed to funding support and manuscript review. H.W. and M.P. provided data for Norway spruce and black spruce, respectively, as well as manuscript review.

## Notes

### Competing Interest Statement

The authors have declared no competing interest.

