## Supplementary Materiel for "Optimizing marker density for maximizing the accuracy of genomic prediction and heritability estimates in three major North American and European spruce species"

Figures S1, S2, and S3 illustrate the relationships existing between prediction accuracy (*PACC*) estimates and the number of SNPs for each trait in each species. Similar to the *PA* results, the findings showed an increase in prediction accuracy with the increasing number of sampled SNPs. This increase stops and stabilizes between 5,000 and 8,000 SNPs. A similar trend is observed between the predictive accuracy of genetic values and breeding values. A slight decrease associated with certain sampling was observed in the datasets with 3,000 and 4,000 SNPs.

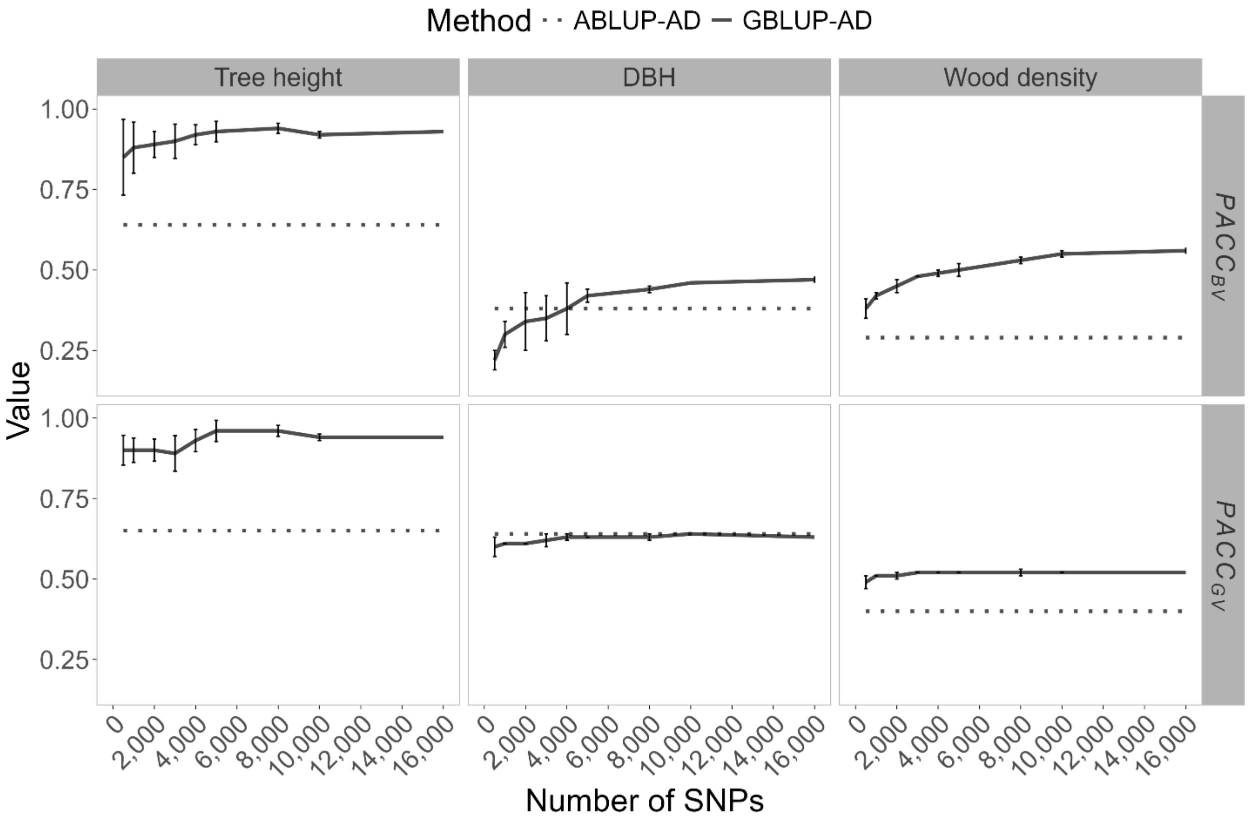


*Fig. S1.* ***Trends of average prediction accuracy of breeding values ($PACC_{BV}$) and genetic values ($PACC_{GV}$) in black spruce GS models (GBLUP-AD) considering varying numbers of SNPs.*** *For the three phenotypic traits the means and standard deviation are shown for ten-fold cross validation of ten times resampled marker subsets. Horizontal dashed lines represent the corresponding pedigree-based models (ABLUP-AD), independent of SNP number. DBH means diameter at breast height.*

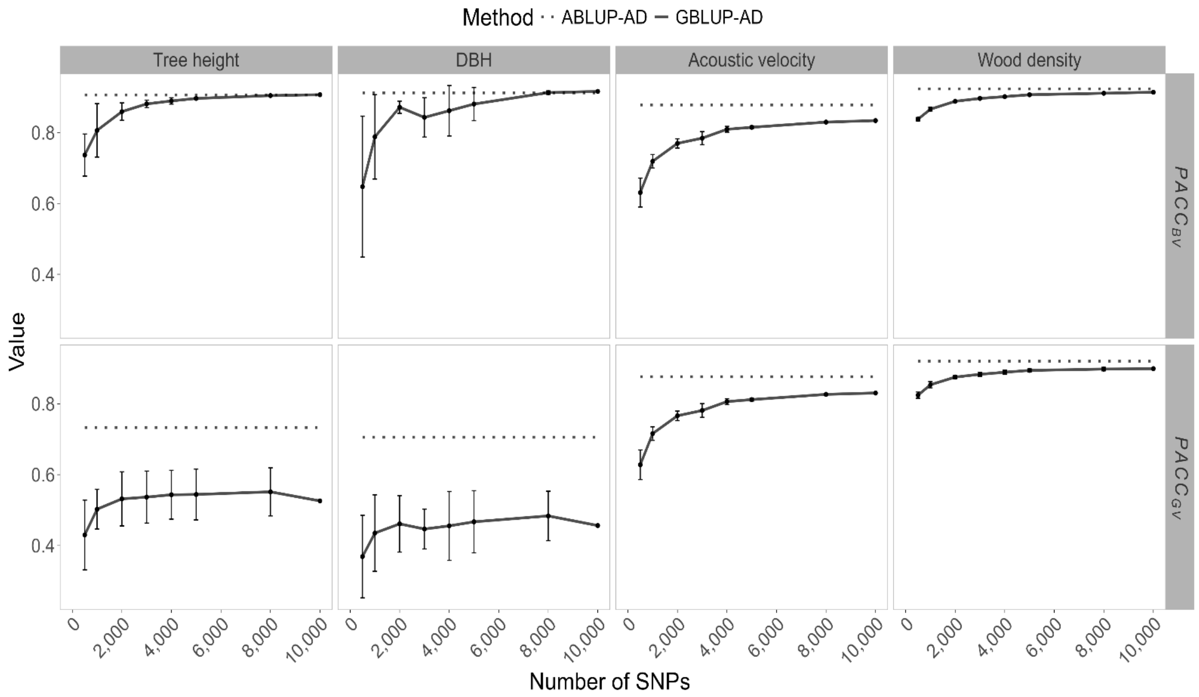


*Fig. S2.* ***Trends of average prediction accuracy of breeding values ($PACC_{BV}$) and genetic values ($PACC_{GV}$) in white spruce GS models (GBLUP-AD) considering varying numbers of SNPs.*** *For the three phenotypic traits the means and standard deviation are shown for ten-fold cross validation of ten times resampled marker subsets. Horizontal dashed lines represent the corresponding pedigree-based models (ABLUP-AD), independent of SNP number. DBH means diameter at breast height.*

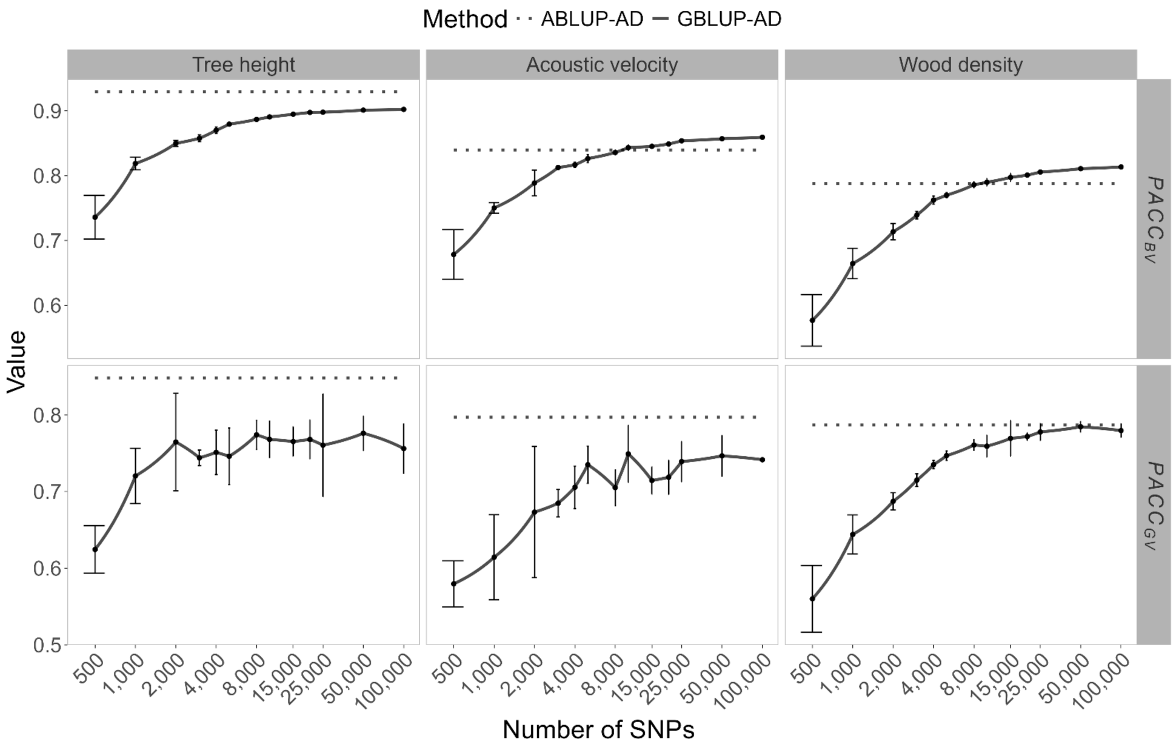


*Fig. S3.* ***Trends of average prediction accuracy of breeding values ($PACC_{BV}$) and genetic values ($PACC_{GV}$) in Norway spruce GS models (GBLUP-AD) considering varying numbers of SNPs.*** *For the three phenotypic traits the means and standard deviation are shown for ten-fold cross validation of ten times resampled marker subsets. Horizontal dashed lines represent the corresponding pedigree-based models (ABLUP-AD), independent of SNP number. The number of SNPs is presented on a logarithmic scale to facilitate readability and interpretability of smaller marker subsets.*

To ensure that five resampling were sufficient, additional resampling up to 20 replicates was performed for one growth trait (tree height) and one wood quality trait (wood density) in white and Norway spruce, the species with the lowest and the highest number of available markers (Fig. S4 et S5)

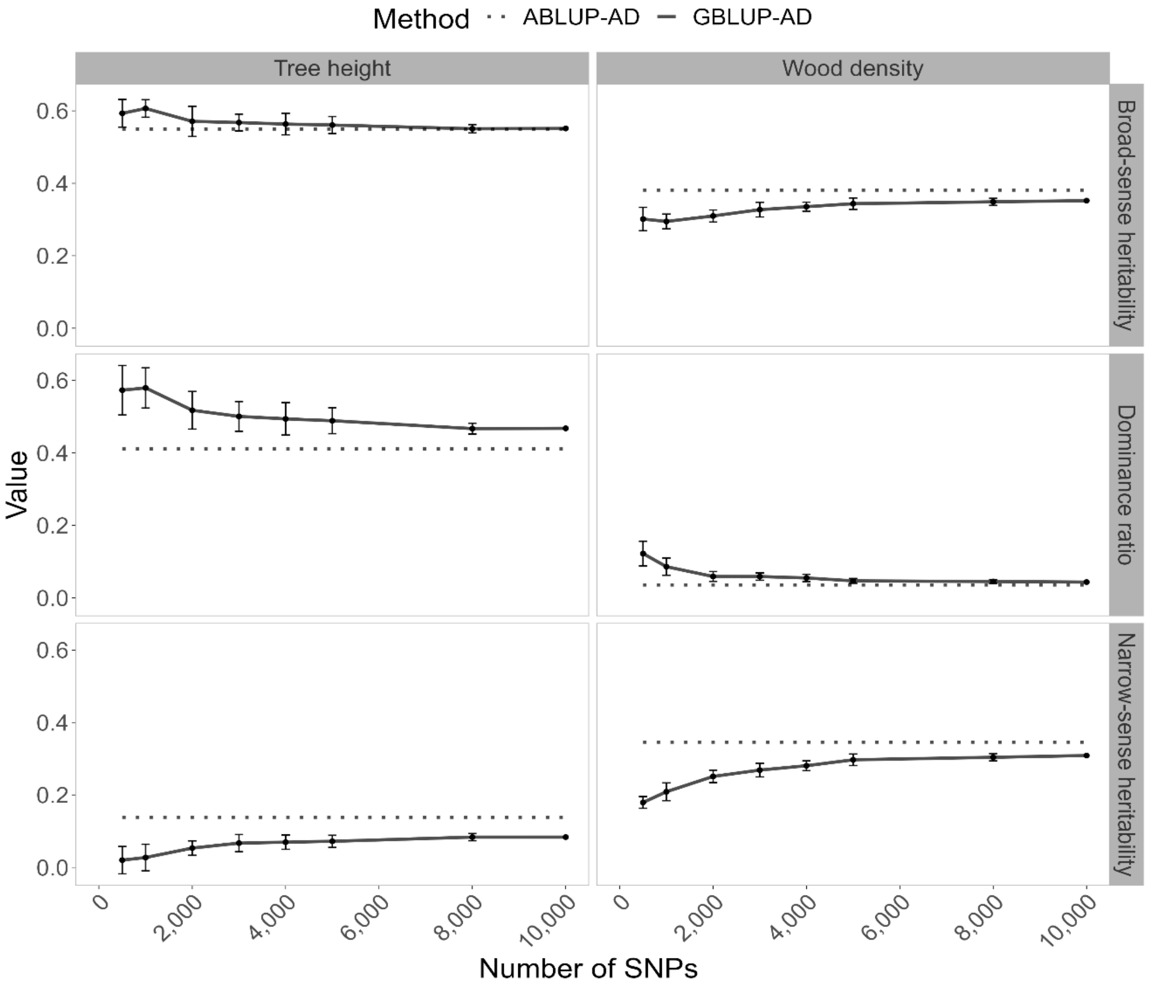


*Fig. S4.* ***White spruce trends in heritability estimates in function of the number of markers used in GS modelling of tree height and wood density****. Broad-sense heritability and its components the narrow-sense heritability and dominance ratio are shown in horizontal panels. Solid lines present the mean estimates of SNP resampling of GBLUP-AD models, including the standard deviations for 20 replicates. The dashed lines represent estimates from the corresponding pedigree-based models (ABLUP-AD).*

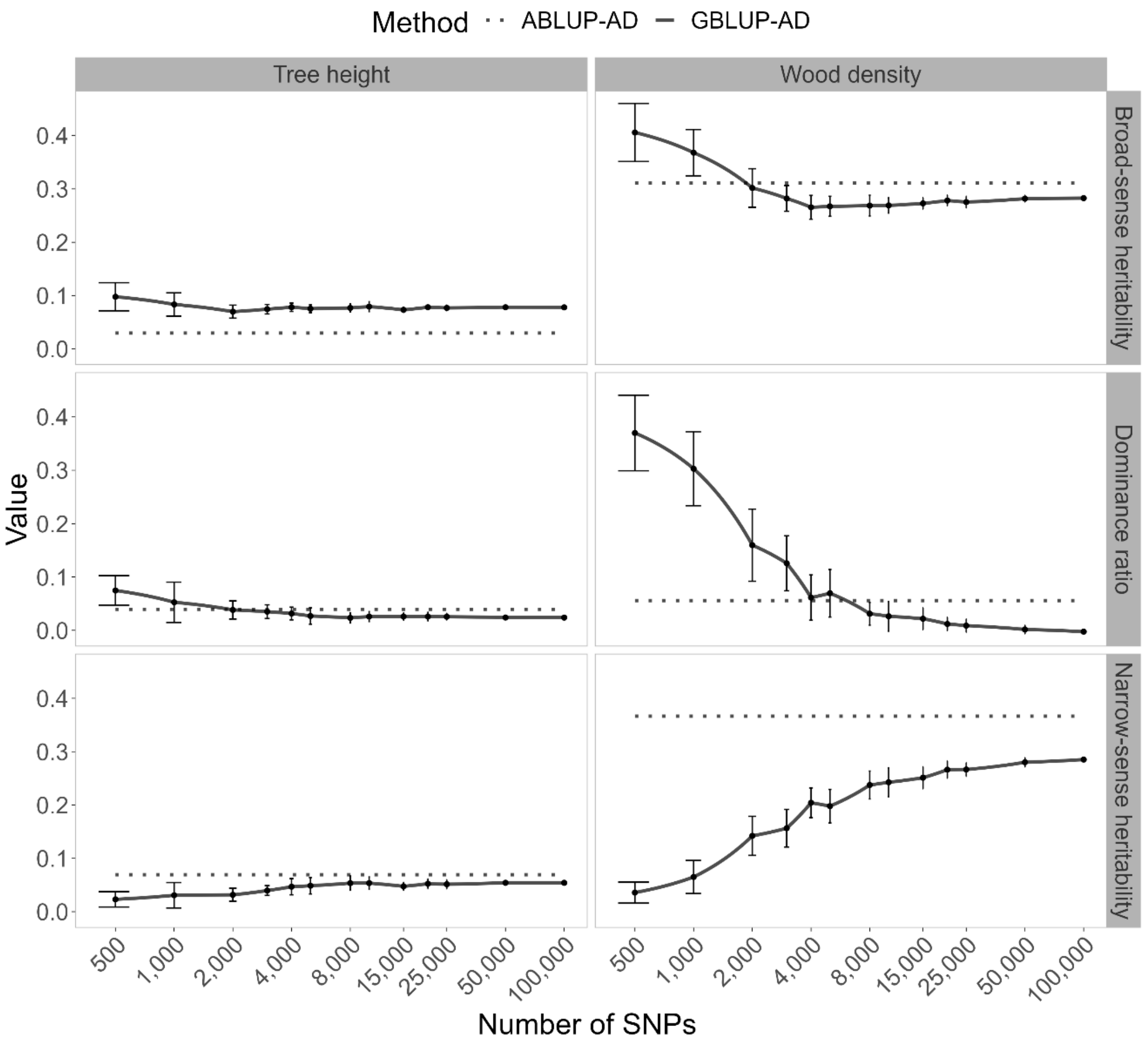


*Fig. S5.* ***Norway spruce trends in heritability estimates in function of the number of markers used in GS modelling of tree height and wood density****. Broad-sense heritability and its components the narrow-sense heritability and dominance ratio are shown in horizontal panels. Solid lines present the mean estimates of SNP resampling of GBLUP-AD models, including the standard deviations for 20 replicates. The dashed lines represent estimates from the corresponding pedigree-based models (ABLUP-AD).*

Figures S6, S7, and S8 present the Spearman correlations of GEBVs, calculated from total genetic values, between each SNP density level and the next higher level. A progressive increase in correlations is observed as the number of SNPs increases.

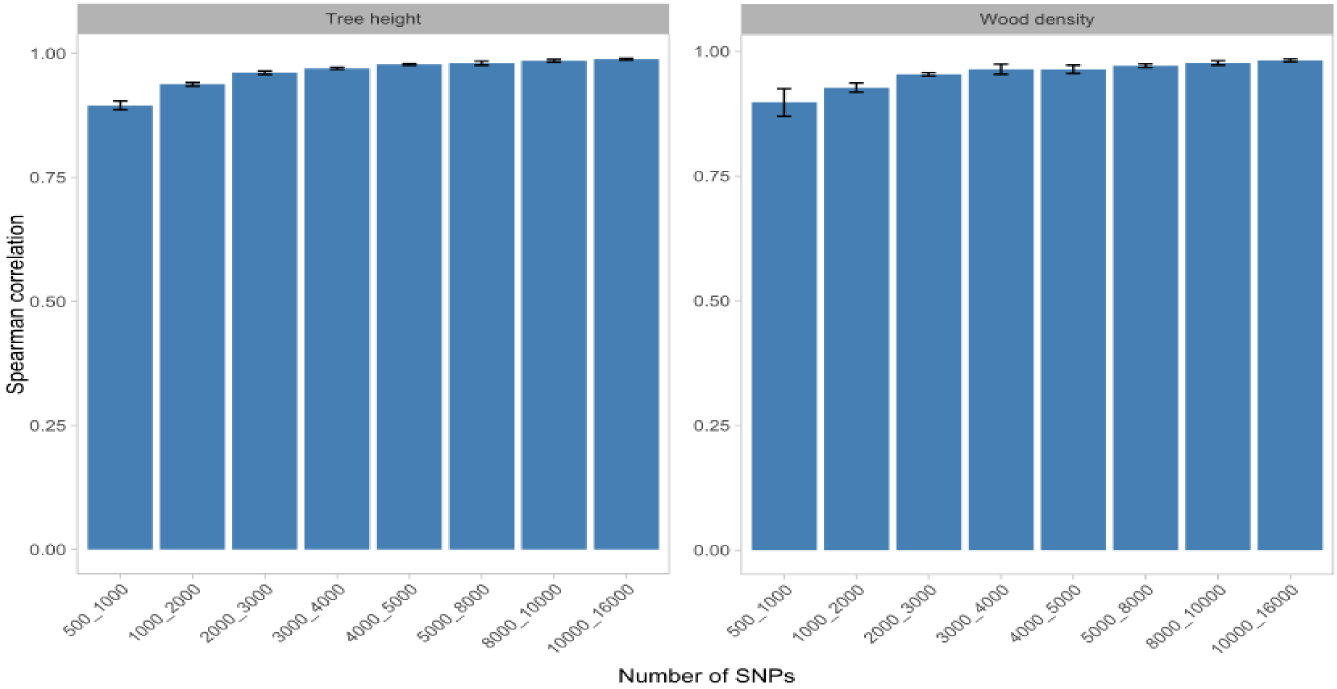


*Fig. S6.* ***Black spruce Spearman correlation trends between successive SNP density levels used in GS modelling of tree height and wood density****. Spearman correlations of GEBVs were calculated based on total genetic values (additive + dominance).*

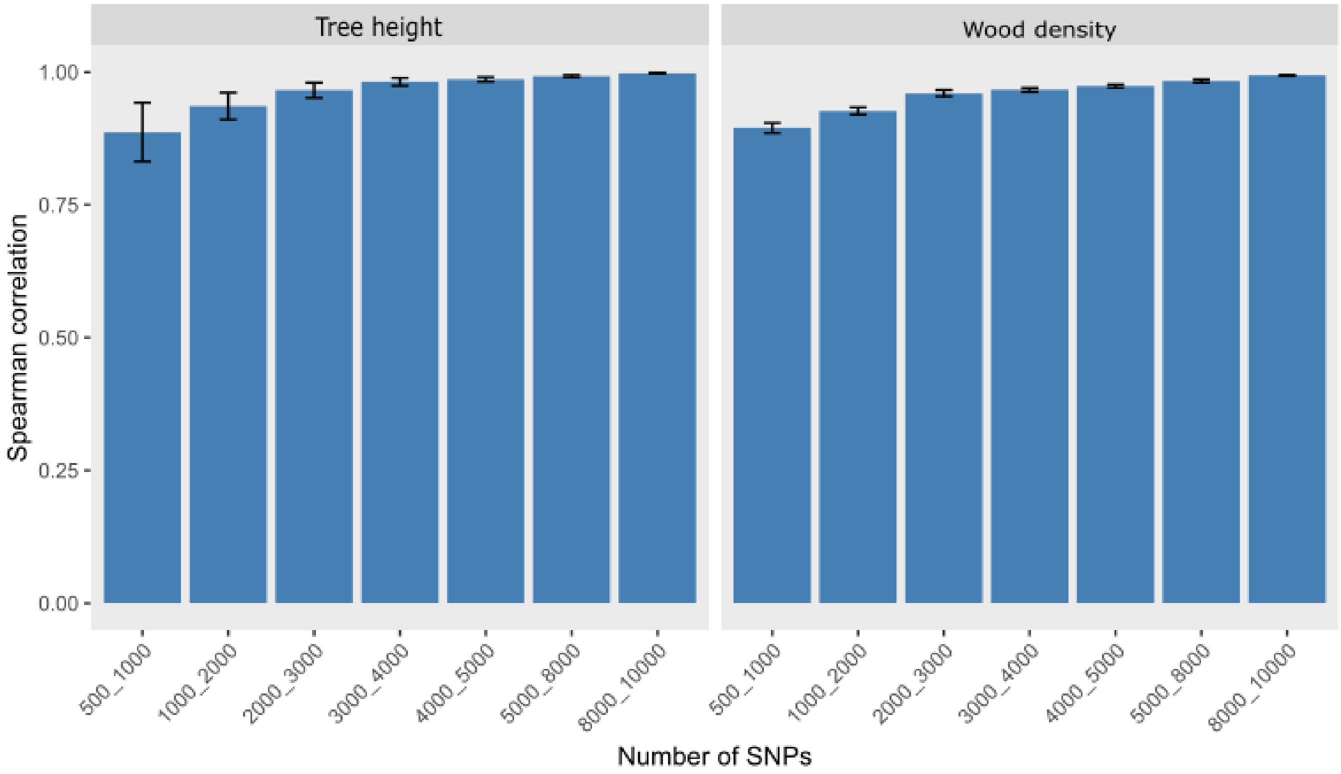


*Fig. S7.* ***White spruce Spearman correlation trends between successive SNP density levels used in GS modelling of tree height and wood density****. Spearman correlations of GEBVs were calculated based on total genetic values (additive + dominance).*

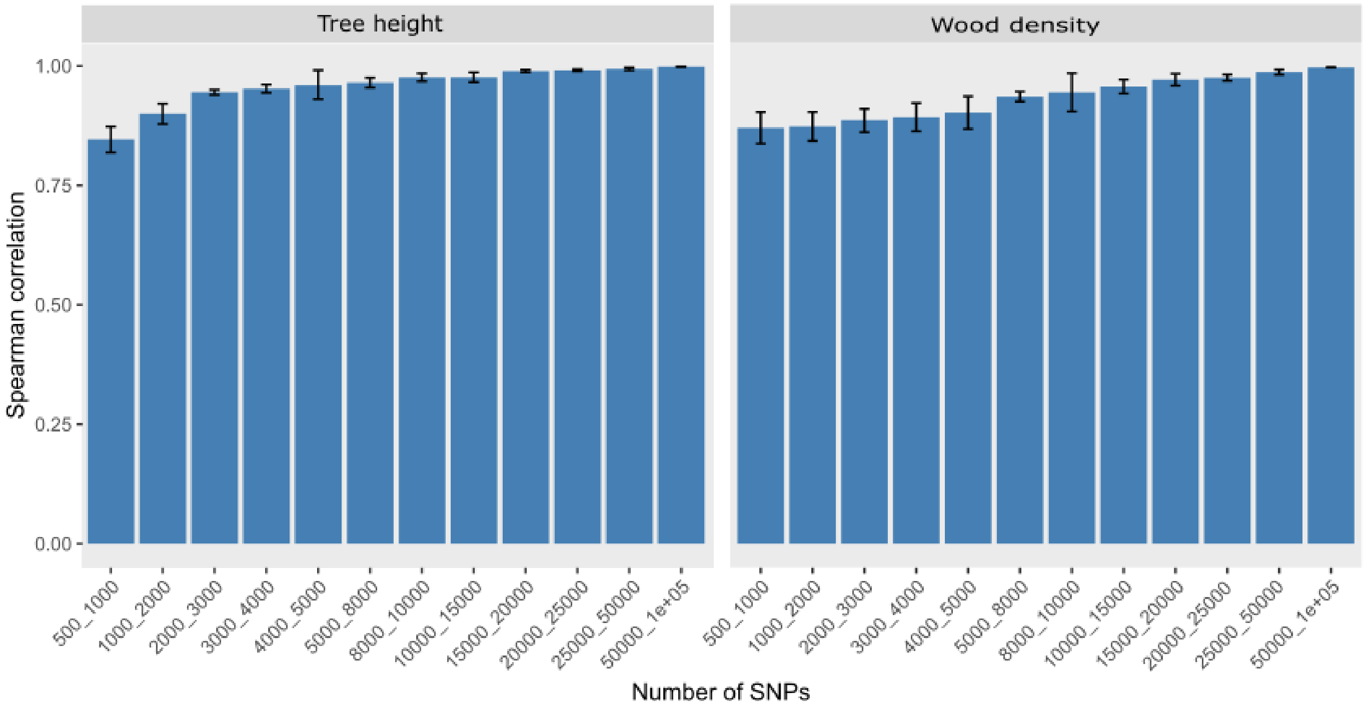


*Fig. S8.* ***Norway spruce Spearman correlation trends between successive SNP density levels used in GS modelling of tree height and wood density****. Spearman correlations of GEBVs were calculated based on total genetic values (additive + dominance).*

**Table S1:** Variance component estimates for black spruce.

| Parameter | Tree height | DBH | Wood density |
|---|---|---|---|
| ABLUP-AD | | | |
| $\hat{\sigma}^2_{b\_MAP}$ | 10.58 (6.10) | 28.60 (20.03) | 34.38 (40.22) |
| $\hat{\sigma}^2_{b\_ROB}$ | 8.57 (5.44) | 20.72 (20.03) | 53.63 (43.01) |
| $\hat{\sigma}^2_{p\_MAP}$ | 35.22 (9.54) | 97.42 (33.95) | 173.45 (180.45) |
| $\hat{\sigma}^2_{p\_ROB}$ | 9.10 (7.42) | 26.31 (23.66) | 168.82 (121.62) |
| $\hat{r}_a$ | 0.89 (0.22) | 0.99 (NA)$^\dagger$ | 0.99 (NA)$^\dagger$ |
| $\hat{\sigma}^2_{a\_MAP}$ | 26.03 (9.75) | 243.11 (103.84) | 976.72 (501.82) |
| $\hat{\sigma}^2_{a\_ROB}$ | 34.15 (8.42) | 261.78 (80.07) | 522.84 (283.21) |
| $\hat{r}_d$ | 0.69 (0.20) | 0.99 (NA)$^\dagger$ | 0.55 (0.28) |
| $\hat{\sigma}^2_{d\_MAP}$ | 0.00 (NA)$^\dagger$ | 169.27 (76.89) | 1159.94 (668.68) |
| $\hat{\sigma}^2_{d\_ROB}$ | 0.00 (NA)$^\dagger$ | 34.10 (44.99) | 788.58 (415.85) |
| $\hat{\sigma}^2_{e\_MAP}$ | 61.33 (9.04) | 0.00 (NA)$^\dagger$ | 41.13 (472.51) |
| $\hat{\sigma}^2_{e\_ROB}$ | 56.23 (8.29) | 63.21 (49.8) | 272.56 (302.64) |
| GBLUP-AD | | | |
| $\hat{\sigma}^2_{b\_MAP}$ | 10.27 (6.79) | 31.62 (21.71) | 32.28 (39.05) |
| $\hat{\sigma}^2_{b\_ROB}$ | 9.06 (5.67) | 21.85 (21.71) | 54.20 (43.40) |
| $\hat{\sigma}^2_{p\_MAP}$ | 35.24 (9.39) | 96.38 (34.11) | 276.92 (175.32) |
| $\hat{\sigma}^2_{p\_ROB}$ | 8.73 (7.26) | 26.28 (24.59) | 177.45 (120.82) |
| $\hat{r}_a$ | 0.91 (0.21) | 0.99 (NA)$^\dagger$ | 0.99 (NA)$^\dagger$ |
| $\hat{\sigma}^2_{a\_MAP}$ | 26.77 (9.30) | 61.87 (30.59) | 343.68 (185.72) |
| $\hat{\sigma}^2_{a\_ROB}$ | 33.40 (7.70) | 60.49 (23.10) | 112.29 (88.12) |
| $\hat{r}_d$ | 0.07 (NA)$^\dagger$ | 0.3 (0.15) | 0.76 (0.14) |
| $\hat{\sigma}^2_{d\_MAP}$ | 0.00 (NA)$^\dagger$ | 300.99 (41.80) | 2135.35 (703.31) |
| $\hat{\sigma}^2_{d\_ROB}$ | 0.00 (NA)$^\dagger$ | 215.95 (84.40) | 1425.02 (171.55) |
| $\hat{\sigma}^2_{e\_MAP}$ | 59.40 (8.87) | 0.00 (NA)$^\dagger$ | 8.76 (541.52) |
| $\hat{\sigma}^2_{e\_ROB}$ | 55.39 (8.06) | 37.59 (67.29) | 0.00 (NA)$^\dagger$ |

[1] DBH is diameter at breast height. Matapedia (MAP), Robidou (ROB) are the sites. $\hat{\sigma}^2_b$ is the block variances; $\hat{\sigma}^2_p$ is the plot variances; $\hat{\sigma}^2_c$ is clonal variance; $\hat{\sigma}^2_a$ and $\hat{\sigma}^2_d$ are the additive and dominance genetic variances respectively; $\hat{r}_a$ and $\hat{r}_d$ the correlation of additive and dominance effects between sites; $\hat{\sigma}^2_e$ are the residual error variances. All models converged. † These terms were at boundary (very close to 0 or 1) after convergence of the model.

**Table S2:** Variance component estimates for white spruce.

| Parameter | Tree height | DBH | Acoustic velocity | Wood density |
|---|---|---|---|---|
| ABLUP-AD | | | | |
| $\hat{\sigma}^2_{b\_ASS}$ | 1438.19 (905.01) | 0.00 (NA)$^{\dagger}$ | 11.54 (8.05) | 52.15 (39.17) |
| $\hat{\sigma}^2_{b\_SCA}$ | 954.68 (696.56) | 20.32 (15.70) | 9.62 (6.82) | 36.47 (28.82) |
| $\hat{\sigma}^2_{p\_ASS}$ | 1045.00 (599.18) | 0.10 (27.99) | 8.23 (8.89) | 105.41 (59.92) |
| $\hat{\sigma}^2_{p\_SCA}$ | 1384.25 (863.59) | 4.15 (28.21) | 11.69 (6.52) | 33.50 (53.13) |
| $\hat{r}_a$ | 0.62 (0.28) | 0.27 (0.42) | 0.93 (0.09) | 1.00 (NA)† |
| $\hat{\sigma}^2_{a\_ASS}$ | 2420.16 (1186.69) | 86.55 (57.32) | 72.90 (23.34) | 362.89 (131.37) |
| $\hat{\sigma}^2_{a\_SCA}$ | 3530.88 (2382.09) | 94.02 (60.95) | 43.69 (16.71) | 257.50 (90.36) |
| $\hat{r}_d$ | 0.99 (NA)$^{\dagger}$ | 0.79 (0.29) | 0.99 (NA)$^{\dagger}$ | 0.99 (NA)† |
| $\hat{\sigma}^2_{d\_ASS}$ | 2015.19 (1776.59) | 196.55 (102.71) | 0.00 (NA)$^{\dagger}$ | 64.64 (130.41) |
| $\hat{\sigma}^2_{d\_SCA}$ | 9023.54 (4191.72) | 181.17 (102.99) | 4.26 (15.54) | 0.00 (NA)† |
| $\hat{\sigma}^2_{e\_ASS}$ | 3824.59 (1374.49) | 165.25 (75.73) | 62.82 (15.59) | 340.00 (119.19) |
| $\hat{\sigma}^2_{e\_SCA}$ | 1163.97 (2882.35) | 183.65 (75.43) | 43.77 (13.41) | 540.16 (79.43) |
| GBLUP-AD | | | | |
| $\hat{\sigma}^2_{b\_ASS}$ | 1436.63 (897.78) | 0.00 (NA)$^{\dagger}$ | 10.81 (7.61) | 57.98 (42.21) |
| $\hat{\sigma}^2_{b\_SCA}$ | 955.54 (696.84) | 19.99 (15.50) | 9.33 (6.60) | 33.67 (26.95) |
| $\hat{\sigma}^2_{p\_ASS}$ | 805.45 (576.88) | 0.00 (NA)$^{\dagger}$ | 9.69 (8.60) | 91.76 (59.28) |
| $\hat{\sigma}^2_{p\_SCA}$ | 1364.25 (857.29) | 4.89 (28.26) | 11.09 (6.06) | 34.93 (51.60) |
| $\hat{r}_a$ | 0.51 (0.29) | -0.13 (0.61) | 0.92 (0.08) | 1.00 (NA)$^{\dagger}$ |
| $\hat{\sigma}^2_{a\_ASS}$ | 2076.34 (862.85) | 45.53 (38.58) | 77.61 (16.90) | 305.78 (89.31) |
| $\hat{\sigma}^2_{a\_SCA}$ | 2294.72 (1452.52) | 54.73 (39.56) | 54.30 (12.64) | 237.04 (68.71) |
| $\hat{r}_d$ | 0.99 (NA)$^{\dagger}$ | 0.83 (0.24) | 0.99(NA)$^{\dagger}$ | 0.99 (NA)$^{\dagger}$ |
| $\hat{\sigma}^2_{d\_ASS}$ | 2333.81 (1509.76) | 251.40 (109.66) | 0.00 (NA)$^{\dagger}$ | 75.53 (125.09) |
| $\hat{\sigma}^2_{d\_SCA}$ | 9838.75 (4013.78) | 213.73 (102.56) | 2.08 (13.55) | 0.00 (NA)$^{\dagger}$ |
| $\hat{\sigma}^2_{e\_ASS}$ | 3869.79 (1269.22) | 144.41 (80.93) | 58.79 (11.90) | 372.42 (109.50) |
| $\hat{\sigma}^2_{e\_SCA}$ | 1157.66 (2915.02) | 178.55 (77.82) | 38.06 (11.83) | 537.13 (71.05) |

[1] DBH is diameter at breast height. Asselin (ASS), Saint-Casimir (SCA) are the sites. $\hat{\sigma}^2_b, \hat{\sigma}^2_p, \hat{\sigma}^2_c, \hat{\sigma}^2_a, \hat{\sigma}^2_d$, $\hat{r}_a$, $\hat{r}_d$ and $\hat{\sigma}^2_e$ are defined in Table 2. All models converged. † These terms were at boundary (very close to 0 or 1) after convergence of the model. The variance component estimates for acoustic velocity were multiplied by 1,000 to ensure better proportionality with the other traits.

**Table S3:** Variance component estimates for Norway spruce.

| Parameter | Tree height | Acoustic velocity | Wood density |
|---|---|---|---|
| ABLUP-AD | | | |
| $\hat{\sigma}^2_{p\_HAD}$ | 1863.20 (628.13) | 3.43 (1.74) | 0.63 (0.24) |
| $\hat{\sigma}^2_{p\_VIN}$ | 798.64 (324.55) | 1.95 (1.32) | 0.24 (0.13) |
| $\hat{r}_a$ | 0.68 (0.42) | 0.85 (0.08) | 0.92 (0.09) |
| $\hat{\sigma}^2_{a\_HAD}$ | 1363.65 (739.26) | 32.56 (9.48) | 2.24 (066) |
| $\hat{\sigma}^2_{a\_VIN}$ | 604.77 (391.38) | 36.09 (10.96) | 2.29 (0.73) |
| $\hat{r}_d$ | -0.2 (0.64) | 0.99 (NA)† | -0.99 (NA)† |
| $\hat{\sigma}^2_{d\_HAD}$ | 2908.21 (1448.17) | 5.95 (6.43) | 0.19 (0.39) |
| $\hat{\sigma}^2_{d\_VIN}$ | 569.78 (925.23) | 11.95 (10.33) | 0.45 (0.74) |
| $\hat{\sigma}^2_{e\_HAD}$ | 6335.02 (1210.66) | 34.73 (7.19) | 2.58 (0.48) |
| $\hat{\sigma}^2_{e\_VIN}$ | 4781.79 (862.34) | 26.86 (10.07) | 2.79 (0.74) |
| GBLUP-AD | | | |
| $\hat{\sigma}^2_{p\_HAD}$ | 1812.32 (613.50) | 3.03 (1.62) | 0.68 (0.25) |
| $\hat{\sigma}^2_{p\_VIN}$ | 757.08 (313.25) | 1.48 (1.14) | 0.22 (0.13) |
| $\hat{r}_a$ | 0.46 (0.35) | 0.71 (0.14) | 0.95 (0.12) |
| $\hat{\sigma}^2_{a\_HAD}$ | 1397.74 (747.73) | 21.15 (6.22) | 1.72 (0.44) |
| $\hat{\sigma}^2_{a\_VIN}$ | 885.29 (421.51) | 33.97 (8.39) | 1.59 (0.50) |
| $\hat{r}_d$ | 0.11 (0.53) | 0.99 (NA)† | -0.03 (0.74) |
| $\hat{\sigma}^2_{d\_HAD}$ | 3524.47 (1483.99) | 26.47 (10.94) | 0.51 (0.51) |
| $\hat{\sigma}^2_{d\_VIN}$ | 640.40 (882.77) | 21.32 (11.18) | 0.72 (0.78) |
| $\hat{\sigma}^2_{e\_HAD}$ | 5691.63 (1270.06) | 23.56 (9.05) | 2.55 (0.47) |
| $\hat{\sigma}^2_{e\_VIN}$ | 4476.26 (858.88) | 19.07 (9.83) | 3.02 (0.72) |

[1] Vindeln (VIN) and Hädanberg (HAD) are the sites. $\hat{\sigma}^2_b$, $\hat{\sigma}^2_p$, $\hat{\sigma}^2_c$, $\hat{\sigma}^2_a$, $\hat{\sigma}^2_d$ , $\hat{r}_a$ , $\hat{r}_d$ and $\hat{\sigma}^2_e$ are defined in Table 2. All models converged. † These terms were at boundary (very close to 0 or 1) after convergence of the model. The variance component estimates for acoustic velocity were multiplied by 1,000 to ensure better proportionality with the other traits.

**Table S4:** Family structure of trees by species.

| | Parents | Family | Tree | Tree means per family | Tree median per family |
|---|---|---|---|---|---|
| Black spruce | 46 | 62 | 912 | 15 | 16 |
| White Spruce | 39 | 56 | 988 | 18 | 18 |
| Norway Spruce | 55 | 127 | 1370 | 10.8 | 10 |
